# Methionine cycle in a pair of serotonergic neurons regulates diet-dependent behavior and longevity through a neuron-gut signaling

**DOI:** 10.1101/2024.03.01.582891

**Authors:** Sabnam Sahin Rahman, Shreya Bhattacharjee, Govind Prakash, Simran Motwani, Tripti Nair, Rachamadugu Sai Keerthana, Arnab Mukhopadhyay

## Abstract

The folate-methionine cycle (Met-C) is a central metabolic pathway that is regulated by vitamin B12 (B12), a micronutrient obtained exclusively from diet and microbiota. This metabolic hub supports amino acid, nucleotide and lipid biosynthesis apart from its central role of providing one carbon (-CH_3_) moiety for methylation reactions. While deficiency of B12 as well as polymorphism in enzymes of the Met-C has been clinically attributed to neurological and metabolic disorders, how this pathway cell non-autonomously regulates systemic physiological processes is less understood. Using a B12-sensitive mutant of *Caenorhabditis elegans*, we show that the neuronal Met-C responds to differential B12 content in diet to regulate p38-MAPK activation in intestinal cells, thereby modulating cytoprotective gene expression, stress tolerance and longevity. Mechanistically, B12-driven changes in the metabolic flux through the Met-C in the serotonergic ADF neurons of the mutant lead to the release of serotonin (5-hydroxytryptamine, 5-HT). 5-HT activates its receptor, MOD-1, in the post-synaptic interneurons that then secretes the neuropeptide FLR-2. FLR-2 binds to FSHR-1, its cognate receptor in the intestine, and induces the phase transition of the SARM domain protein TIR-1, thereby activating the p38-MAPK pathway. Importantly, this cascade influences the foraging behaviour of the mutant worms such that they prefer a B12-rich diet. Together, our study reveals a dynamic neuron-gut signaling axis that helps an organism modulate behaviour and life history traits based on the neuronal Met-C metabolic flux determined by B12 availability in its diet. Understandably, disruption of the optimum functioning of this axis may have debilitating effects on the health of an organism and the survival of the species.

## Introduction

Diet typically comprises macronutrients like carbohydrates, fats, and proteins, but also serves as a vital source of essential micronutrients like B12. Notably, an organism is entirely dependent on dietary sources and its microbiota for the supply of B12 (Watanabe 2007). B12 exists in two biologically active forms, methylcobalamin and adenosylcobalamin, functioning as coenzymes for methionine synthase and methylmalonyl-CoA mutase, respectively. The former is required in the folate-methionine cycle (Met-C) of the one-carbon metabolism, facilitating the transfer of methyl group from methyltetrahydrofolate to homocysteine, resulting in the formation of methionine **(Figure 1A)**. Methionine, in conjunction with ATP, undergoes further conversion to S-adenosylmethionine (SAM). SAM, in turn, serves as the primary methyl donor for the methylation of DNA, RNA or other cellular metabolites, and is converted to S-adenosyl homocysteine (SAH). Met-C also supplies key metabolites for the biosynthesis of amino acids, nucleotides and lipids (Parkhitko et al. 2019). Thus, B12 deficiency leads to increased susceptibility to disorders like depression, anxiety, and schizophrenia (Tufan et al. 2012; Mitchell, Conus, and Kaput 2014; Sangle et al. 2020; Zhang et al. 2016) while polymorphism in the genes of the Met-C is linked to increased risk of obesity, insulin resistance, cardiovascular disease and sarcopenia (Kim et al. 2010; Lisboa et al. 2020; Di Renzo et al. 2019). However, the precise mechanisms underpinning the association between B12 deficiency, Met-C polymorphism and their pathological outcomes are not fully understood. Importantly, whether and how B12-driven changes in Met-C in one tissue may affect systemic physiology through effects on a distal tissue is less known.

**Figure 1.**
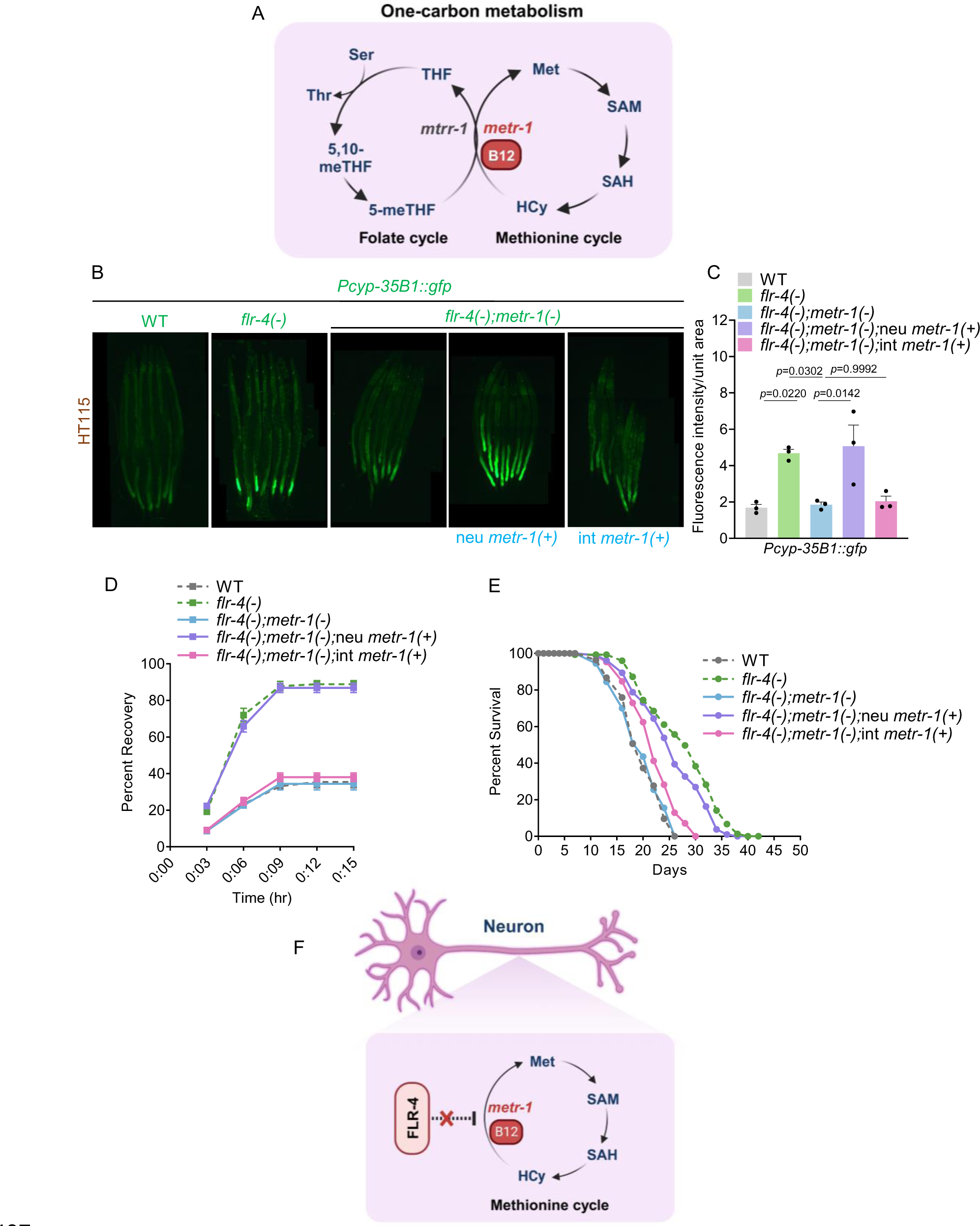
Met-C in the neuron modulates CyTP gene expression, osmotic tolerance and life span of *flr-4(n2259)*. (A) A schematic representation of one-carbon metabolism in *Caenorhabditis elegans*, consisting of the folate and methionine cycle (Met-C). (B) The expression of *gfp* in *flr-4(n2259);metr-1(ok521);Pcyp35B1::gfp* worms was restored when *metr-1* was rescued only in the neurons (using the pan-neuronal *rgef-1* promoter) [neu *metr-1(+)*] but not when rescued in the intestine (using the *ges-1* promoter) [int *metr-1(+)*]. One of three biologically independent replicates is shown. (C) Quantification of (B). Average of three biological replicates ± SEM. *P*-value determined using One-way ANOVA with Tukey’s multiple comparisons test. (D) The osmotic tolerance of *flr-4(n2259);metr-1(ok521)* worms were restored when *metr-1* was rescued only in the neurons (using the pan-neuronal *rgef-1* promoter) [neu *metr-1(+)*] but not when rescued in the intestine (using the *ges-1* promoter) [int *metr-1(+)*]. One of three biologically independent replicates is shown. (E) The life span of *flr-4(n2259);metr-1(ok521)* worms were restored when *metr-1* was rescued only in the neurons (using the pan-neuronal *rgef-1* promoter) [neu *metr-1(+)*] but not when rescued in the intestine (using the *ges-1* promoter) [int *metr-1(+)*]. One of two biologically independent replicates is shown. (F) A schematic illustrating that the Met-C is required in the neurons for *flr-4(-)* phenotypes. Summary of osmotic stress tolerance and life span assays are provided in the Source Data file. All experiments were performed at 20 °C. All data and analysis are provided in the Source Data file.

Invertebrate model systems, like *Caenorhabditis elegans*, are playing a prominent role in our efforts to understand the contributions of B12 and Met-C in cellular physiology, highlighting the importance of host-microbiota interactions on a spectrum of life-history traits, gene expression dynamics, metabolic adaptation, behavior and neuronal health (Wei and Ruvkun 2020; Giese et al. 2020; Zhang et al. 2016; Na et al. 2018; Kenyon 2005; Kenyon et al. 1993; MacNeil et al. 2013; Yilmaz and Walhout 2014). *C. elegans* is a bacterivore that feeds on diverse bacterial species in both its natural habitats and in controlled laboratory settings; thus, bacteria act both as food and the microbiota of the worms. So, like mammals, the worms are entirely dependent on their bacterial diet for B12 supply. Additionally, due to the availability of diverse genetic, biochemical and cell biology tools, it serves as a popular model system for the exploration of evolutionarily conserved genetic pathways regulating cellular stress responses, diseases as well as inter-tissue crosstalk (Van Pelt and Truttmann 2020; Rodriguez et al. 2013; Durieux, Wolff, and Dillin 2011; Taylor and Dillin 2013; Berendzen et al. 2016; Bar-Ziv et al. 2023; Miller et al. 2022; Burkewitz et al. 2015; Miller et al. 2020).

In multicellular organisms, inter-tissue communications are important for maintaining physiological homeostasis. To study the inter-tissue crosstalk that mediates the effect of B12 through the modulation of the Met-C, a sensitized genetic background is necessary. This is because the wild-type worms that feed on a wide range of bacterial diets, can maintain cellular and metabolic homeostasis orchestrated by specific genes, preventing alteration of life-history traits on different diets rich in diverse nutrients. However, occasionally we come across genetic mutants that show altered life-history traits specific to one diet and not others. These ‘gene-diet pairs’ have significantly contributed to our knowledge of the impact of food quality and microbiota on both life span and health (Yen and Curran 2016). Our laboratory has characterized one such diet-gene pair where a serine-threonine kinase gene mutant displays an increased sensitivity to B12 and exhibits enhanced cytoprotective gene (CyTP) expression, stress tolerance and life span only on the B12-rich *E. coli* HT115 but not on *E. coli* OP50 (Verma et al. 2018), providing us with the perfect paradigm to study the tissue-specific regulation of Met-C by B12 and its effect on distal tissues.

When the kinase-dead allele *flr-4(n2259)* [referred to as *flr-4(-)*] is fed HT115 or OP50 supplemented with B12, the metabolic flux through the Met-C cycle is augmented, leading to the downstream activation of the p38-MAPK pathway (Nair et al. 2022). Notably, *flr-4* expresses both in the neurons as well as the intestine and knocking down the gene using RNAi in either tissue partially increases life span (Verma et al. 2018), suggesting that the gene may independently affect the Met-C and the p38-MAPK pathway in distinct tissues. Here, we show that the B12-driven changes in Met-C metabolic flux takes place in the serotonergic ADF neurons while the p38-MAPK is activated in the intestine. 5-HT secreted from these neurons binds to its cognate receptor MOD-1 in the post-synaptic interneuron which then releases the glycoprotein hormone/neuropeptide FLR-2. FLR-2 binds to its receptor in the intestinal cells and directs the phase transition of the SARM domain-containing TIR-1 protein that activates the p38 MAPK pathway. This results in the increase in CyTP expression, stress tolerance and longevity. Importantly, this axis also regulates the foraging behaviour of the mutant, directing them towards the B12-rich diet. Together, our study elucidates a novel neuron-gut signaling cascade that transmits the information of neuronal Met-C to modulate gene expression in the distal intestinal cells to regulate life span in response to a diet of varying B12 content.

## Results

### The neuronal Met-C is essential for CyTP expression, stress tolerance, and longevity of the *flr-4* mutant

In our previous study, we demonstrated that the *flr-4(-)* worms exhibit increased health span and life span when grown on a B12-rich diet such as HT115 or OP50 supplemented with B12. This effect is attributed to the increased flux through the Met-C (Nair et al. 2022). B12 acts as a cofactor for the evolutionarily conserved methionine synthase (METR-1) enzyme, facilitating the conversion of homocysteine to methionine **(Figure 1A)**.

Since the *flr-4* expression is primarily confined to neuronal and intestinal cells (Verma et al. 2018), we asked in which specific tissue would *flr-4* interact with the Met-C to regulate downstream processes to ensure longevity. To discern this, we restored the expression of the *metr-1* cDNA selectively in either the neuronal cells of *flr-4(-);metr-1(-)*, using the pan-neuronal *rgef-1* promoter *[neu metr-1(+)],* or intestinal cells, using the *ges-1* promoter *[int metr-1(+)].* We used the transgenic strain *Pcyp35B1::gfp*, where the promoter of a CyTP gene (*cyp35B1*) drives the expression of *gfp*, that faithfully reports p38-MAPK activation in *flr-4(-)* worms in response to B12 (Verma et al. 2018; Nair et al. 2022). It may be noted that the increased CyTP gene expression, stress tolerance and life span of *flr-4(-)* is suppressed when *metr-1* is knocked down using RNAi (Nair et al. 2022) or knocked out using *metr-1* null mutant [*flr-4(-);metr-1(-)*] (**Figure 1 B-E)**. We found that *flr-4(-);metr-1(-);Prgef-1::metr-1;Pcyp35B1::gfp* worms when grown on HT115, where *metr-1* is rescued in the neuronal cells *[neu metr-1(+)]*, exhibited increased expression of CyTP genes compared to control worms (**Figure 1B,C**). However, *flr-4(-);metr-1(-);Pges-1::metr-1,Pcyp35B1::gfp* worms, where *metr-1* was rescued in the intestinal cells *[int metr-1(+)]*, did not display enhanced GFP expression. Similarly, neuronal restoration of *metr-1* enhanced the osmotic stress tolerance (**Figure 1D**) as well as the lifespan (**Figure 1E**) of *flr-4(-);metr-1(-)* mutant worms to the level observed in *flr-4(-)*. We could attribute these benefits to the high B12 levels in HT115 (Nair et al. 2022) as supplementing the OP50 with B12 could also restore the CyTP gene expression (**Figure S1A,B**) as well as osmotic stress tolerance (**Figure S1C,D,E**) only in the neuronally-expressed *metr-1* strain. Collectively, these observations provide evidence that the Met cycle is crucial in the neurons, but not in the intestine, for the lifespan extending phenotypic traits observed in the *flr-4(-)* worms (**Figure 1F**).

### p38-MAPK signaling in the intestine activates CyTP gene expression, stress tolerance and longevity of the *flr-4* mutant

The *flr-4(-)* worms, grown on a B12-rich diet, exhibit an increased life span and health span attributed to the activation of the p38-MAPK pathway (Verma et al. 2018; Nair et al. 2022). In *C. elegans,* this pathway acts as a central signaling mediator required for mounting an innate immune response when challenged with pathogens. The worm p38 ortholog PMK-1 is activated by its upstream MAPKKK NSY-1 and MAPKK SEK-1 (**Figure 2A**) (Kim et al. 2002). We aimed to delineate the specific tissue types where the NSY-1-SEK-1-PMK-1 signaling module is crucial for *flr-4(-)* to manifest the benefits. To accomplish this, we used a tissue-specific RNAi system, selectively knocking down *sek-1* either in the neurons or in the intestine. The CyTP gene expression was suppressed when *sek-1* was knocked down in *flr-4(n2259);rde-1(ne219);Pnhx-::rde-1;Pcyp35B1::gfp* [intestine (int) only RNAi], while it remained unaffected in *flr-4(n2259);sid-1(pk3321);Punc-119::sid-1;Pcyp35B1::gfp* [neuron (neu) only RNAi] worms (**Figure 2B,C**). To further validate the involvement of intestinal *sek-1,* we generated an *flr-4(-);sek-1(-);Pges-1::sek-1* strain that has *sek-1* cDNA expressed only in the intestinal cells [int *sek-1(+)*] and *flr-4(-);sek-1(-);Punc-119::sek-1* strain that expresses *sek-1* only in the neuronal cells [neu *sek-1(+)*]. We found that only the intestinal restoration of *sek-1* enhanced the osmotic stress tolerance (**Figure 2D**) as well as the longevity (**Figure 2E**) of *flr-4(-);sek-1(-)* worms to that of *flr-4(-)* worms. Collectively, these observations provide evidence that the p38-MAPK pathway is required in the intestine for the phenotypic traits observed in the *flr-4(-)* mutants (**Figure 2F**).

**Figure 2.**
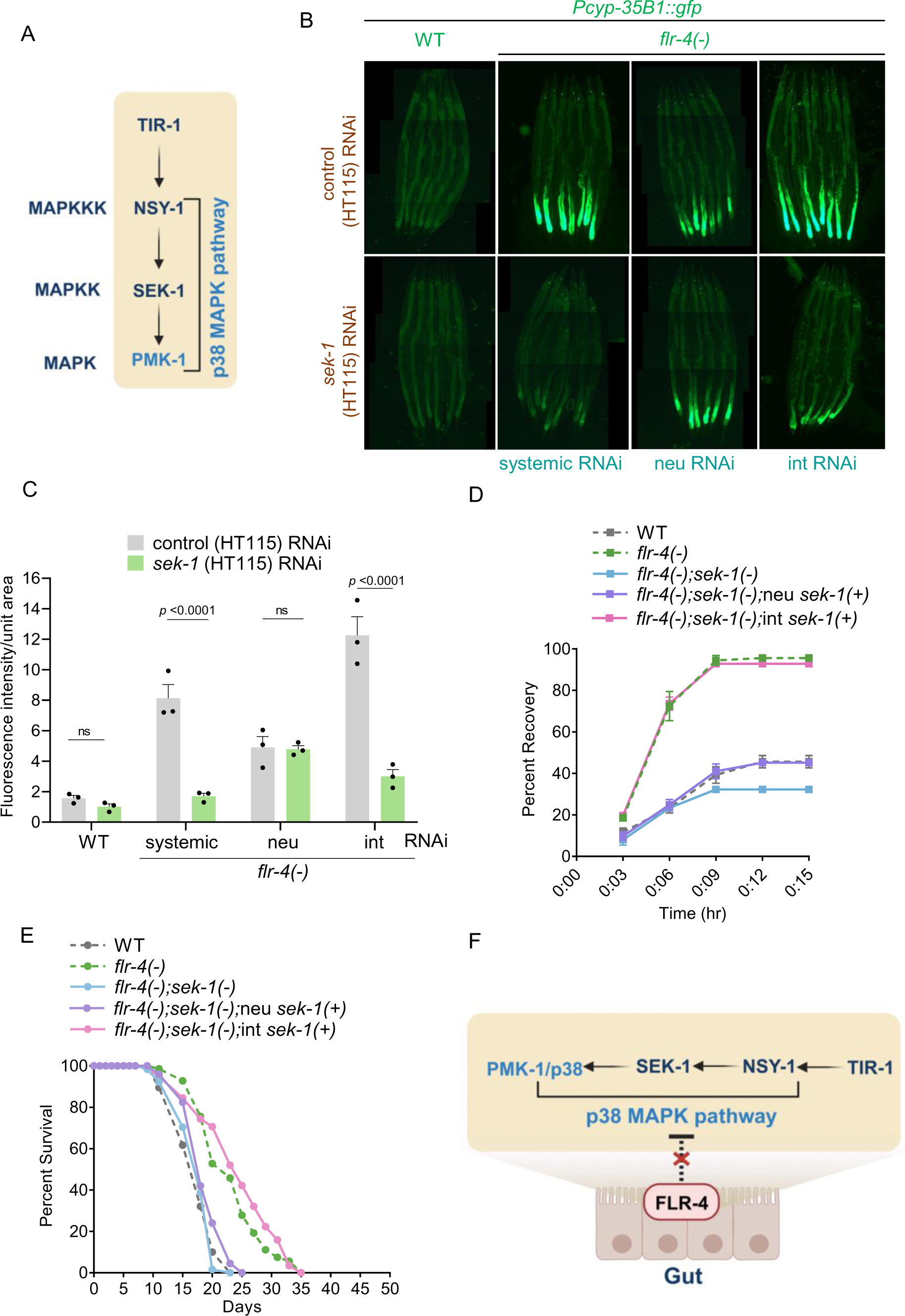
p38-MAPK pathway in the intestine modulates CyTP gene expression, osmotic tolerance and life span of *flr-4(n2259)*. (A) A schematic representation of the p38-MAPK pathway in *C. elegans*. (B) The expression of *gfp* was suppressed when *sek-1* was knocked down in *flr-4(n2259);rde-1(ne219);Pnhx-2::rde-1;Pcyp35B1::gfp* [intestine only RNAi (int RNAi)] but not in *flr-4(n2259);sid-1(pk3321);Punc-119::sid-1;Pcyp35B1::gfp* [neuron only RNAi (neu RNAi)] worms. One of three biologically independent replicates is shown. (C) Quantification of (B). Average of three biological replicates ± SEM. *P*-value determined using Two-way ANOVA with Tukey’s Multiple Comparison Test. (D) The osmotic tolerance was restored when *sek-1* was rescued only in the intestine of the *flr-4(n2259);sek-1(km4)* worms (using the *ges-1* promoter) [int *sek-1(+)*] but not when rescued in the neurons (using the pan-neuronal *unc-119* promoter) [neu *sek-1(+)*]. One of three biologically independent replicates is shown. (E) The life span was restored when *sek-1* was rescued only in the intestine of the *flr-4(n2259);sek-1(km4)* worms (using the *ges-1* promoter) [int *sek-1(+)*] but not when rescued in the neurons (using the pan-neuronal *unc-119* promoter) [neu *sek-1(+)*]. One of three biologically independent replicates shown. (F) A schematic illustrating that the p38-MAPK pathway is required in the gut. Summary of osmotic stress tolerance and life span assays are provided in the Source Data file. All experiments were performed at 20 °C. All data and analysis are provided in the Source Data file.

### The CyTP gene expression, stress tolerance and longevity of the *flr-4* mutant require serotonergic neurotransmission

Since the Met-C is crucial in the neurons and the p38-MAPK pathway in the intestine govern CyTP gene expression, stress tolerance and longevity phenotypes in the *flr-4(-)* mutant worms, we investigated how this inter-tissue communication is established (**Figure 3A**). The dense core vesicles are required for neuropeptide release while the clear synaptic vesicles are involved in neurotransmitter release (Miller et al. 1996; Cornell et al. 2022). We used *flr-4(-);Pcyp31B1::gfp* worms and knocked down the representative genes required for neurotransmitter release (*unc-13*) or neuropeptide release (*unc-31*) using RNAi. Interestingly, we observed that in either of the scenarios the increased CyTP gene expression was diminished, suggesting that *flr-4(-)* engages certain downstream neuropeptides and neurotransmitters (**Figure S2A,B**). To identify the neurotransmitters, we systemically knocked down genes involved in dopamine (*cat-2*), glutamate (*eat-4*), octopamine (*tbh-1*), tyramine (*tdc-1*), serotonin (*tph-1*), acetylcholine (*unc-17*), or GABA (*unc-25*) biosynthesis in the *flr-4(-);Pcyp31B1::gfp* using RNAi. We found that the expression of *gfp* was suppressed on knocking down *tph-1* and *cat-2*, highlighting the importance of serotonergic and dopaminergic neurotransmission in the *flr-4(-)* phenotype (**Figure S2C**). For further characterization of the *flr-4(-)* signaling cascade, in this study, we only report the involvement of the serotonergic arm.

**Figure 3.**
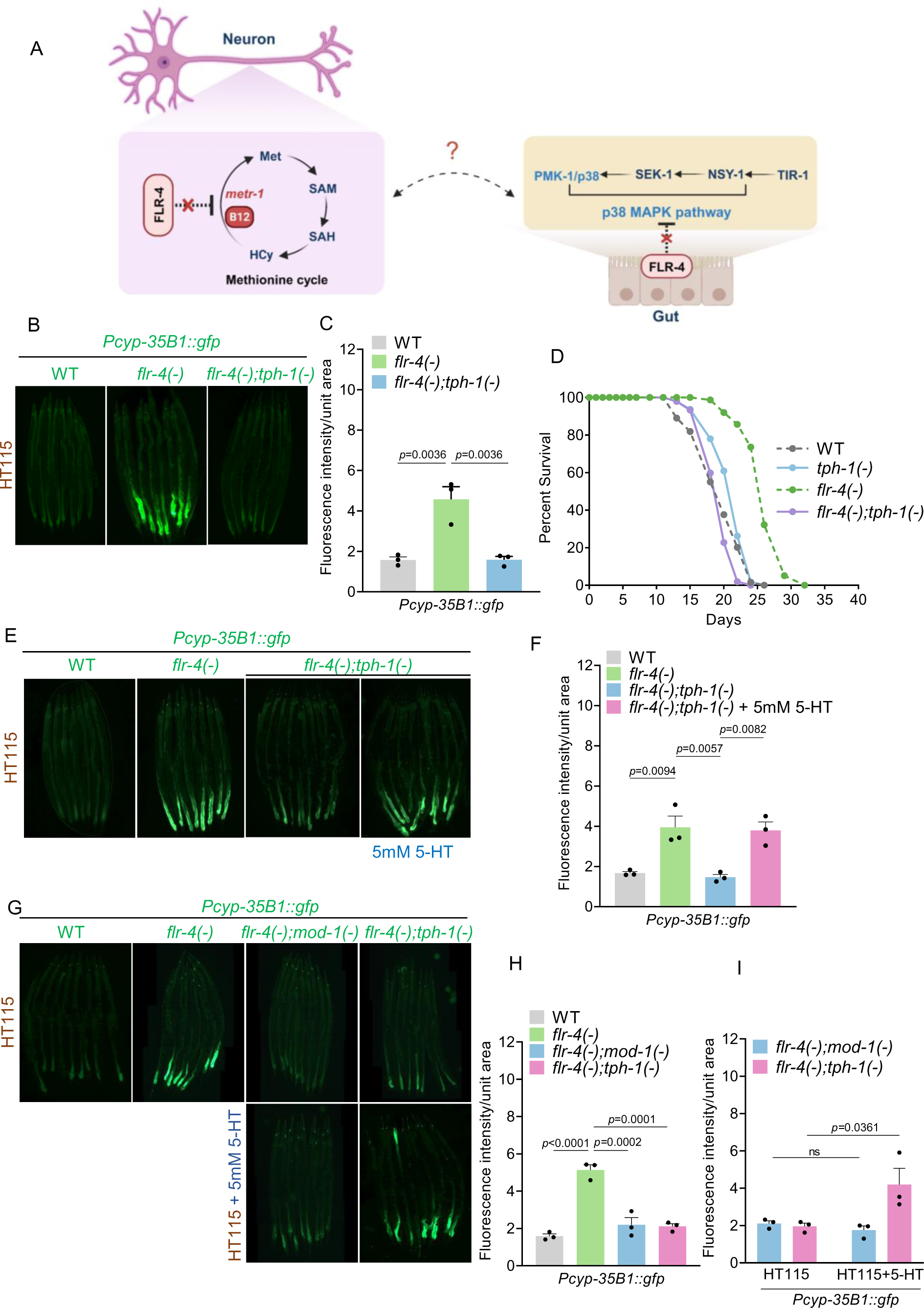
The activation of CyTP gene expression and longevity of *flr-4(n2259)* worms requires serotonergic signaling. (A) A schematic illustrating the potential inter-tissue crosstalk of the neuron-gut axis. (B) The expression of *gfp* was suppressed in case of *flr-4(n2259);tph-1(mg280);Pcyp35B1::gfp* worms compared to *flr-4(n2259);Pcyp35B1::gfp* worms. One of three biologically independent replicates is shown. (C) Quantification of (B). Average of three biological replicates ± SEM. *P*-value determined using One-way ANOVA with Tukey’s multiple comparisons test. (D) The increased life span of *flr-4(n2259)* worms was suppressed when *tph-1* was mutated as in *flr-4(n2259);tph-1(mg280)* worms. One of three biologically independent replicates is shown. (E) The expression of *gfp* was restored when *flr-4(n2259);tph-1(mg280);Pcyp35B1::gfp* worms were supplemented with 5mM 5-HT. One of three biologically independent replicates is shown. (F) Quantification of (E). Average of three biological replicates ± SEM. *P*-value determined using One-way ANOVA with Tukey’s multiple comparisons test. (G) The expression of *gfp* in *flr-4(n2259);Pcyp35B1::gfp* worms was suppressed when *mod-1* was mutated as in *flr-4(n2259);mod-1(ok103);Pcyp35B1::gfp.* The supplementation of 5mM 5-HT failed to restore *gfp* expression in *flr-4(n2259);mod-1(ok103);Pcyp35B1::gfp* but not in *flr-4(n2259);tph-1(mg280);Pcyp35B1::gfp* worms. One of three biologically independent replicates is shown. (H-I) Quantification of (G). Average of three biological replicates ± SEM. *P*-value determined using One-way ANOVA with Tukey’s multiple comparisons test (G) and Two-way ANOVA with Tukey’s multiple comparisons test (H). Summary of life span assays are provided in the Source Data file. All experiments were performed at 20 °C. All data and analysis are provided in the Source Data file.

We validated the involvement of serotonergic signaling in *flr-4(-)* phenotypes using a genetic deletion of *tph-1* gene (codes for tryptophan hydroxylase, the enzyme that catalyzes the first step of serotonin biosynthesis) as in *flr-4(-);tph-1(-);Pcyp-35B1::gfp*; the expression of the CyTP reporter was suppressed (**Figure 3B,C**). We also found that the increased life span of *flr-4(-)* is completely suppressed by *tph-1* deletion (**Figure 3D**). A recent study has established the role of phenylalanine hydroxylase (*pah-1)* in the biosynthesis of non-neuronal 5-HT (Yu et al. 2023). So, to further investigate serotonergic neurotransmission, we first determined whether non-neuronal 5-HT was also involved. For this, we knocked down *pah-1* and found that this did not affect the expression of *gfp* in *flr-4(-);Pcyp35B1::gfp*, pointing to the specific involvement of the neuronally synthesized 5-HT in *flr-4(-)* phenotype (**Figure S3A,B**). To confirm the role of 5-HT, we supplemented the neurotransmitter to the *flr-4(-);tph-1(-);Pcyp-35B1::gfp* worms where the *gfp* expression is suppressed due to the lack of serotonergic signaling. We found that the expression of *gfp* was restored (**Figure 3E,F**). However, supplementation of 5-HT to the *Pcyp-35B1::gfp* and *flr-4(-);Pcyp-35B1::gfp* worms did not affect the CyTP gene expression (**Figure S3C**).

5-HT functions by binding to its cognate receptor at the postsynaptic neurons. To identify the 5-HT receptors that act downstream of *flr-4*, we made double mutants of *flr-4(-);Pcyp-35B1::gfp* with each of the four receptor mutants, i.e., *ser-1(-)*, *ser-4(-)*, *ser-7(-),* and *mod-1(-)* (Carre-Pierrat et al. 2006). We found that *mod-1* mutation significantly suppressed the CyTP gene expression of the *flr-4(-);Pcyp-35B1::gfp* worms (**Figure S3D,E**) along with their osmotic tolerance (**Figure S3F**) and longevity (**Figure S3G**). Importantly, supplementing 5-HT to *flr-4(-);mod-1(-);Pcyp35B1::gfp* failed to increase *gfp* expression, suggesting that MOD-1 is the cognate receptor of 5-HT essential for downstream effects of *flr-4(-)* (**Figure 3G,H,I**).

### Serotonergic neurotransmission connects neuronal Met-C to the intestinal p38-MAPK pathway

Till now, we have only provided evidence of the requirement of serotonergic neurotransmission downstream of *flr-4*. So next, to determine the hierarchy of the serotonergic neurotransmission in the Met-C-p38-MAPK cascade downstream of *flr-4*, we used supplementation assays. First, we supplemented 5-HT to *flr-4(-);metr-1(-);Pcyp35B1::gfp* worms, where *gfp* fluorescence is low, and found that the expression was restored (**Figure 4A,B**). Similarly, supplementation of 5-HT to *flr-4(-);metr-1(-)* worms increased the osmotic tolerance to the extent seen in *flr-4(-)* worms (**Figure S4A**).

**Figure 4.**
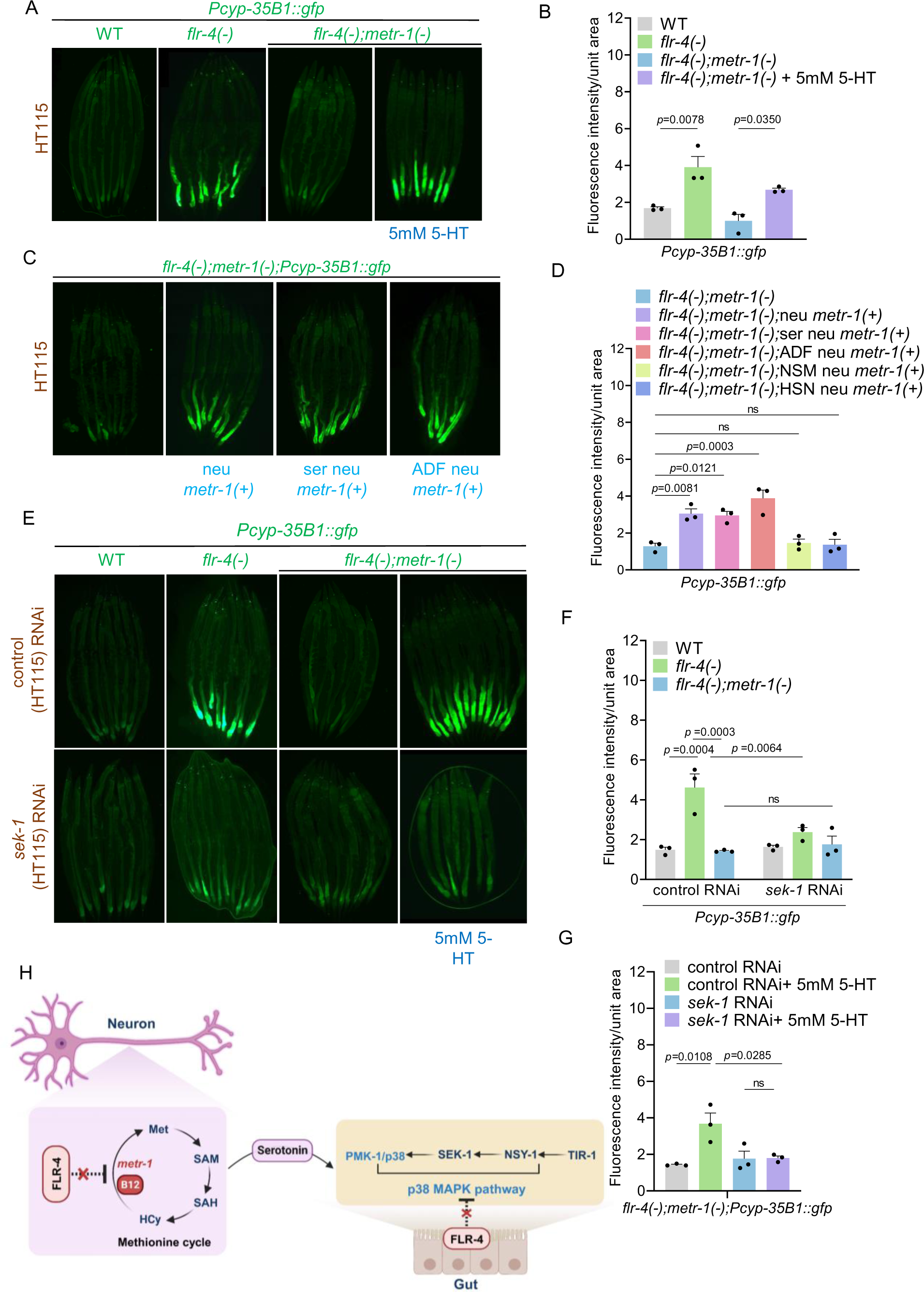
Serotonergic neurotransmission connects the neuronal Met-C to the p38-MAPK pathway in the intestine. (A) The expression of *gfp* in *flr-4(n2259);metr-1(ok521);Pcyp35B1::gfp* worms was restored when these worms were supplemented with 5mM 5-HT. One of three biologically independent replicates is shown. (B) Quantification of (A). Average of three biological replicates ± SEM. *P*-value determined using One-way ANOVA with Tukey’s multiple comparisons test. (C) The expression of *gfp* in *flr-4(n2259);metr-1(ok521);Pcyp35B1::gfp* worms was restored when *metr-1* was rescued only in the serotonergic neurons (using the *tph-1* promoter) [*ser neu metr-1(+)*] or in the ADF serotonergic neurons (using the *srh-142* promoter) [ADF *neu metr-1(+)*]. One of three biologically independent replicates is shown. (D) Quantification of (C) and (S4B). Average of three biological replicates ± SEM. *P*-value determined using One-way ANOVA with Tukey’s multiple comparisons test. (E) The expression of *gfp* in *flr-4(n2259);metr-1(ok521);Pcyp35B1::gfp* worms failed to increase on supplementation with 5mM 5-HT when *sek-1* was knocked down by RNAi. One of three biologically independent replicates is shown. (F-G) Quantification of (E). Average of three biological replicates ± SEM. *P*-value determined using Two-way ANOVA with Tukey’s multiple comparisons test (F) and One-way ANOVA with Tukey’s multiple comparisons test (G). (H) A schematic illustrating the modulation of neuron-gut crosstalk through serotonergic signaling. All experiments were performed at 20 °C. All data and analysis are provided in the Source Data file.

To ascertain the adequacy of rescuing the Met-C exclusively within the serotonergic neurons (*ser neu*) of *flr-4(-)* worms for the faithful recapitulation of the observed phenotypes, we transgenically rescued the *metr-1* cDNA using the *tph-1* promoter. Interestingly, we observed that reconstituting the Met-C specifically within the serotonergic neurons, using the *tph-1* promoter to drive expression of *metr-1*, was sufficient to increase the CyTP gene expression to the levels similar to worms where the pan-neuronal *rgef-1* promoter was used (**Figure 4C,D**) and enhance the osmotic stress tolerance (**Figure S4C**) of the *flr-4(-);metr-1(-)* worms.

Next, we identified the particular serotonergic neurons involved in modulating the Met-C. *C elegans* has three pairs of serotonergic neurons; the chemosensory ADF neurons that perceive external environmental signals (Bargmann and Horvitz 1991; Zhang, Lu, and Bargmann 2005; Song et al. 2013), the pharyngeal secretory NSM neurons that respond to food stimuli (Rhoades et al. 2019) and the egg-laying HSN motor neurons responsible for vulval contraction (Lloret-Fernandez et al. 2018). We rescued *metr-1* cDNA within the ADF neurons (using the *srh-142* promoter), the NSM neuron (using the *tph-1* short promoter), and within the HSN neurons (using the *egl-6* promoter) (Flavell et al. 2013; Rhoades et al. 2019). Met-C restoration specifically in the ADF neurons was sufficient to increase the CyTP gene expression (**Figure 4C,D,S4B**) and osmotic stress tolerance of the *flr-4(-);metr-1(-)* worms (**Figure S4C**). Collectively, these observations strongly indicate that the Met-C exerts its influence within the ADF serotonergic neurons leading to increased serotonergic neurotransmission.

It is also technically possible that serotonergic signaling may work downstream of the p38-MAPK to activate the CyTP genes. In that case, 5-HT supplementation should be independent of a functional p38-MAPK pathway. We tested this by supplementing 5-HT to the *flr-4(-);metr-1(-);Pcyp-35B1::gfp* worms grown on control or *sek-1* RNAi. We found that while the expression of *gfp* was restored on control RNAi, but on *sek-1* RNAi, the expression failed to increase (**Figure 4E,F,G**). Additionally, the increased osmotic tolerance observed in *flr-4(-);metr-1(-)* when supplemented with 5-HT, was not noticed in the *flr-4(-);sek-1(-)* worms (**Figure S4D**), proving that the p38-MAPK signaling works downstream of the serotonergic signaling, connecting Met-C status of the serotonergic neurons to p38-MAPK activation in the intestine (**Figure 4H**).

### FLR-2/FSHR-1 signaling functions downstream of the serotonergic signaling

While the serotonergic signaling connects the neuronal Met-C to intestinal p38-MAPK activation in the *flr-4(-)* mutant, the 5-HT receptor MOD-1 that we identified expresses in the interneurons and not in the intestine (Zhang, Lu, and Bargmann 2005; Churgin et al. 2017; Ranganathan, Cannon, and Horvitz 2000). In a screen aimed at identifying fluoride resistance (*flr)* genes, the *flr-4* was originally identified as one of the class 1 genes (*flr-1, flr-3* and *flr-4*). These mutants exhibited a strong temperature-sensitive defecation defect. The screen also identified the presence of the *flr-2* which belongs to the class 2 *flr* genes (*flr-2, flr-5, flr-6* and *flr-7*), whose mutations suppress certain phenotypes associated with the class 1 genes (Katsura et al. 1994; Iwasaki, Liu, and Thomas 1995; Oishi et al. 2009). FLR-2 is expressed in neurons and encodes a secreted protein that shares significant similarity with the human glycoprotein hormone subunit α2 of thyrostimulin (Oishi et al. 2009; Park et al. 2005). So, we asked whether the neuropeptide FLR-2 could connect the signals from the MOD-1-expressing interneurons to the intestine.

In this direction, we had previously shown that knocking down *unc-31*, a gene which is involved in neuropeptide release from dense-core vesicles, suppresses the expression of *gfp* in *flr-4(-);Pcyp35B1::gfp* (**Figure S2A,B**). This prompted us to check the involvement of FLR-2 neuropeptide in modulating the *flr-4(-)* phenotype, connecting the MOD-1-containing interneuron to the intestine (**Figure 5A**). We found that the *flr-2* mutation suppressed the *gfp* expression of *flr-4(-);Pcyp35B1::gfp* worms (**Figure 5B,C**). Additionally, the osmotic stress tolerance (**Figure 5E**) and the life span (**Figure 5G**) of *flr-4(-)* worms were suppressed when *flr-2* was mutated. FLR-2 appears to function downstream of the 5-HT signaling as 5-HT supplementation could not restore the expression of *gfp* in *flr-4(-);flr-2(-);Pcyp35B1::gfp* worms or the osmotic tolerance of the *flr-4(-);flr-2(-)* worms (**Figure 5B,C,D,F**).

**Figure 5.**
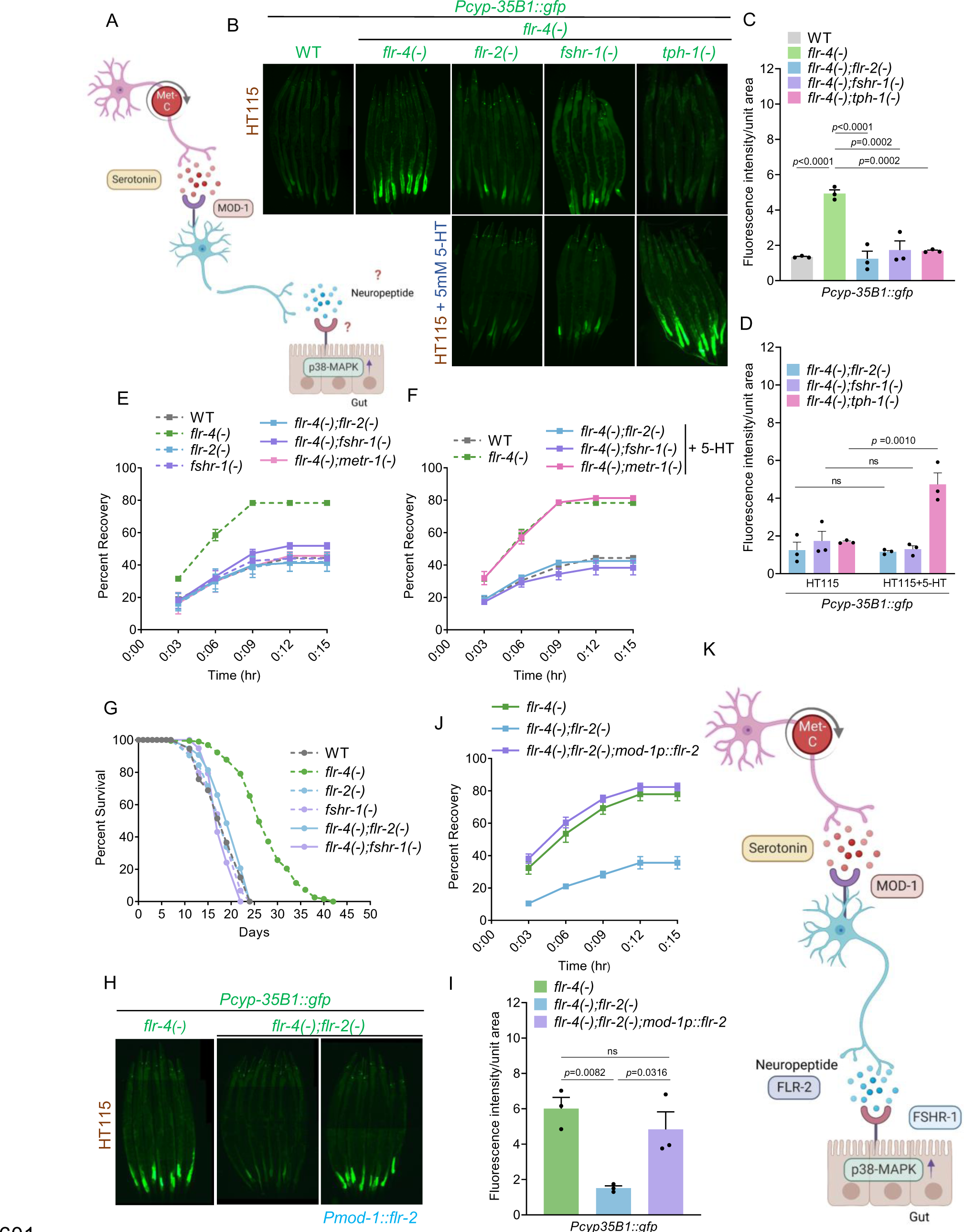
Involvement of the FLR-2/FSHR-1 axis in CyTP gene expression, osmotic tolerance, and life span of *flr-4(n2259)*. (A) A schematic diagram showing the involvement of a potential neuropeptide signaling acting downstream of the Met-C and serotonin. (B) The expression of *gfp* in *flr-4(n2259);Pcyp35B1::gfp* worms was suppressed when neuropeptide *flr-2* or its receptor *fshr-1* was mutated, as in *flr-4(n2259);flr-2(ut5);Pcyp35B1::gfp or flr-4(n2259);fshr-1(ok778);Pcyp35B1::gfp*, respectively. Additionally, supplementation with 5mM 5-HT failed to restore the *gfp* expression in either strain. One of three biologically independent replicates is shown. (C-D) Quantification of (B). Average of three biological replicates ± SEM. *P*-value determined using One-way ANOVA with Tukey’s multiple comparisons test (C) and Two-way ANOVA with Tukey’s multiple comparisons test (D). (E) Osmotic tolerance of *flr-4(n2259)* worms was suppressed when neuropeptide *flr-2* or its receptor *fshr-1* was mutated, as in *flr-4(n2259);flr-2(ut5) or flr-4(n2259);fshr-1(ok778)*, respectively. One of three biologically independent replicates is shown. (F) Supplementation of 5mM 5-HT to *flr-4(n2259);flr-2(ut5) or flr-4(n2259);fshr-1(ok778)* failed to restore osmotic tolerance. One of three biologically independent replicates is shown. (G) The life span of *flr-4(n2259)* worms was suppressed when neuropeptide *flr-2* or its receptor *fshr-1* was mutated, as in *flr-4(n2259);flr-2(ut5) or flr-4(n2259);fshr-1(ok778)*, respectively. One of two biologically independent replicates is shown. (H) The expression of *gfp* in *flr-4(n2259);flr-2(ut5);Pcyp35B1::gfp* worms was restored when neuropeptide *flr-2* cDNA was driven by the *mod-1* promoter as in *flr-4(n2259);flr-2(ut5);mod-1p::flr-2;Pcyp35B1::gfp* worms. One of three biologically independent replicates is shown. (I) Quantification of (H). Average of three biological replicates ± SEM. *P*-value determined using One-way ANOVA with Tukey’s multiple comparisons test. (J) Osmotic stress tolerance of *flr-4(n2259);flr-2(ut5)* worms was restored when neuropeptide *flr-2* cDNA was driven by the *mod-1* promoter as in *flr-4(n2259);flr-2(ut5);mod-1p::flr-2* worms. One of three biologically independent replicates is shown. (K) Model establishing FLR-2 as the neuropeptide secreted by MOD-1 expressing interneurons and FSHR-1 as its intestinal receptor, downstream of the serotonergic signaling pathway. Summary of osmotic stress tolerance and life span assays are provided in the Source Data file. All experiments were performed at 20 °C. All data and analysis are provided in the Source Data file.

Having established that FLR-2 functions downstream of the serotonergic signaling, we then asked whether the neurons expressing MOD-1 are indeed the ones responsible for the release of FLR-2, thereby transducing signals from the serotonergic neurons to the intestine. Interestingly, upon transgenically expressing *flr-2* cDNA under the control of the *mod-1* promoter, we observed increased expression of the CyTP gene (**Figure 5H,I**) and enhanced tolerance to osmotic stress (**Figure 5J**). Thus, MOD-1-expressing neurons serve as the recipient for serotonergic signals and in turn, release the neuropeptide FLR-2 for downstream activation of the p38 MAPK pathway in the intestine.

Next, we wanted to identify the cognate receptor for FLR-2 that transmits signals to the intestine. The intestinally-expressed FSHR-1 is a known receptor for FLR-2, facilitating signal transduction in response to cold-induced stress (Wang et al. 2023) and also to developmental cues (Torzone et al. 2023). To determine whether FSHR-1 also works downstream of the FLR-4 cascade, we visualized the expression of *gfp* in the *flr-4(-);fshr-1(-);Pcyp35B1::gfp* worms. We observed that the expression of *gfp* was reduced, showing that FSHR-1 functions in this pathway (**Figure 5B,C**). Evidently, the osmotic stress tolerance (**Figure 5E**) and the life span (**Figure 5G**) of *flr-4(-)* worms was suppressed when *fshr-1* was mutated. Importantly, like FLR-2, FSHR-1 also acts downstream of the 5-HT signaling as 5-HT supplementation could not restore the expression of *gfp* (**Figure 5B,C,D**) in *flr-4(-);fshr-1(-);Pcyp35B1::gfp* worms or the osmotic tolerance (**Figure 5F**) of the *flr-4(-);fshr-1(-)* worms.

Next, we validated whether FSHR-1 is required in the intestine to support *flr-4(-)* phenotypes. For this, we specifically knocked down *fshr-1* in the intestine of the *flr-4(-);Pcyp35B1::gfp* or *flr-4(-)*, using an intestine-specific RNAi strain, which suppressed *gfp* expression (**Figure S5A,B**) and osmotic stress tolerance (**Figure S5C**), respectively. Importantly, systemic and intestinal knockdown, but not the neuronal knockdown, of *fshr-1* suppressed the long lifespan of *flr-4(-)* worms (**Figure S5D**). This shows that FLR-4-Met-C-serotonin-FLR-2 acts upstream of the intestinal FSHR-1 to activate the p38-MAPK signaling (**Figure 5K**).

### The Met-C-serotonin-FLR-2/FSHR-1 axis induces the TIR-1/SARM1 phase transition to activate the p38-MAPK pathway

Thus far, we have shown that the Met-C functions in the serotonergic neurons while the p38-MAPK works in the intestine of the *flr-4(-)* worms, connected by the serotonin-FLR-2/FSHR-1 axis. Next, we asked how the binding of FLR-2 on intestinal FSHR-1 receptor activates the p38-MAPK pathway.

The TIR domain adaptor protein TIR-1, an ortholog of human SARM1, acts upstream of the p38-MAPK pathway to modulate the expression of genes associated with immune responses (Couillault et al. 2004; Liberati et al. 2004). The initiation of TIR-1 activation is prompted by a stress-induced phase transition, which facilitates protein oligomerization, augmenting NAD+ glycohydrolase activity, and thereby activating the NSY-1/SEK-1/PMK-1 pathway (Peterson et al. 2022). So, we asked whether TIR-1/SARM1 phase transition is induced in the *flr-4(-)* worms to activate the p38-MAPK pathway. For this, we generated the *flr-4(-);tir-1::wrmScarlet* worms that reports TIR-1 expression (Peterson et al. 2022) and observed a significant increase in the formation of visible puncta of the multimerized TIR-1::wrmScarlet protein within the intestinal epithelial cells of *flr-4(-)* worms, compared to the wild-type worms (**Figure 6A,B**). Next, to investigate whether the Met-C-serotonin-FLR-2/FSHR-1 axis downstream of *flr-4* drives the phase transition of TIR-1/SARM, we generated *flr-4(-);metr-1(-);tir-1::wrmScarlet, flr-4(-);tph-1(-);tir-1::wrmScarlet, flr-4(-);flr-2(-);tir-1::wrmScarlet* and *flr-4(-);fshr-1(-);tir-1::wrmScarlet*. As expected, abrogation of any of the components of the axis in the *flr-4(-)* mutant worms significantly dampened the TIR-1 phase transition (**Figure 6B,S6A**). Thus, *flr-4(-)* worms induce TIR-1 oligomerization through the Met-C-serotonin-FLR-2-FSHR-1 axis, triggering the activation of the p38-MAPK pathway to ensure longevity (**Figure 6C**).

**Figure 6.**
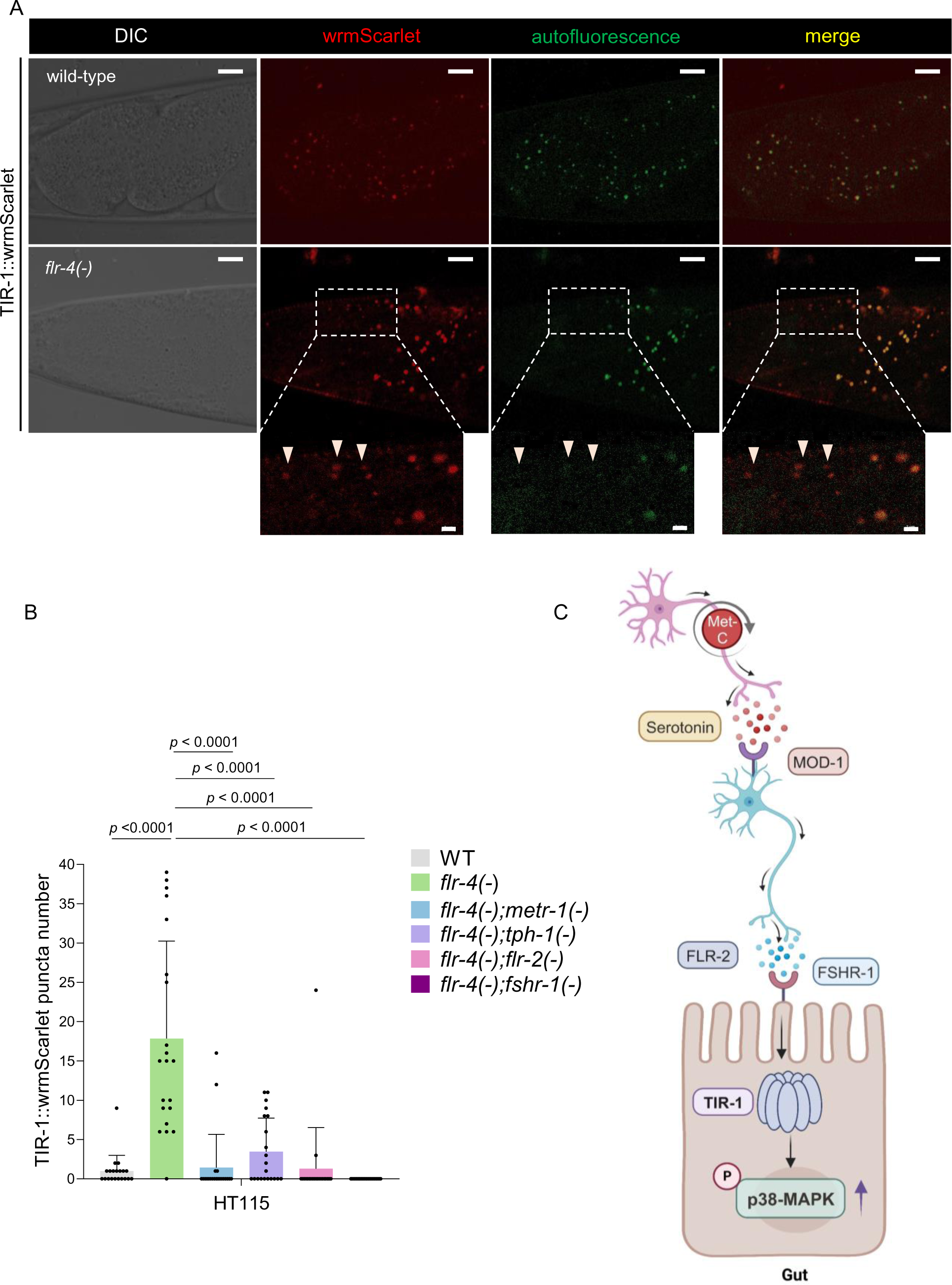
*flr-4(n2259)* worms induce SARM1/TIR-1 phase transition. (A) The *flr-4(n2259)* worms displayed increased TIR-1 puncta formation in comparison to wild-type worms. The red fluorescence channel images captured both TIR-1::wrmScarlet fluorescence and autofluorescence signals, while the green channel images exclusively revealed signals originating from the autofluorescent gut granules (Scale bar 10 μm). TIR-1::wrmScarlet puncta are denoted by arrowheads in the magnified images (Scale bar 2 μm). (B) Quantification of (A) and (S6A). Average of three biological replicates ± SD. *P*-value determined using One-way ANOVA with Tukey’s multiple comparisons test. (C) Model showing TIR-1 oligomerization leading to activation of the p38-MAPK pathway mediated by B12-rich diet. All experiments were performed at 20 °C. All data and analysis are provided in the Source Data file.

### Neuronal Met-C modulates foraging behavior through the serotonin-FLR-2/FSHR-1-p38-MAPK axis

Animals display food preferences or aversions based on environmental cues and nutritional requirements (Rainwater and Guler 2022; Anderson and Li 1987; Johnson 2013). Previous studies have revealed that the worms can make dietary choices based on the nutritional quality of the food as well as their consequent effects on the survival trajectories (Harris et al. 2014; Abada et al. 2009; Shtonda and Avery 2006). Since the *flr-4(-)* worms benefit from the high B12 HT115 diet in terms of enhanced longevity, we asked whether these worms would be naturally inclined to choose HT115 over OP50. To determine this, wild-type and *flr-4(-)* worms were conditioned on either OP50 or HT115 (grown from L1 to L4 stage) and then allowed to choose between these two diets kept on the opposite poles of a 100 mm plate (**Figure 7A**). We observed that while the wild-type worms exhibited no preference towards either diet, interestingly, the *flr-4(-)* mutant worms conditioned on any diet consistently displayed a preference to the HT115 (**Figure 7B**). Next, worms either conditioned on OP50 or OP50 supplemented with B12, were used for the food choice experiment. The *flr-4(-)* worms exhibited a preference towards B12 supplemented OP50 diet, independent of preconditioning, suggesting that these worms were responding specifically to the high B12 levels of HT115 (**Figure S7A**).

**Figure 7.**
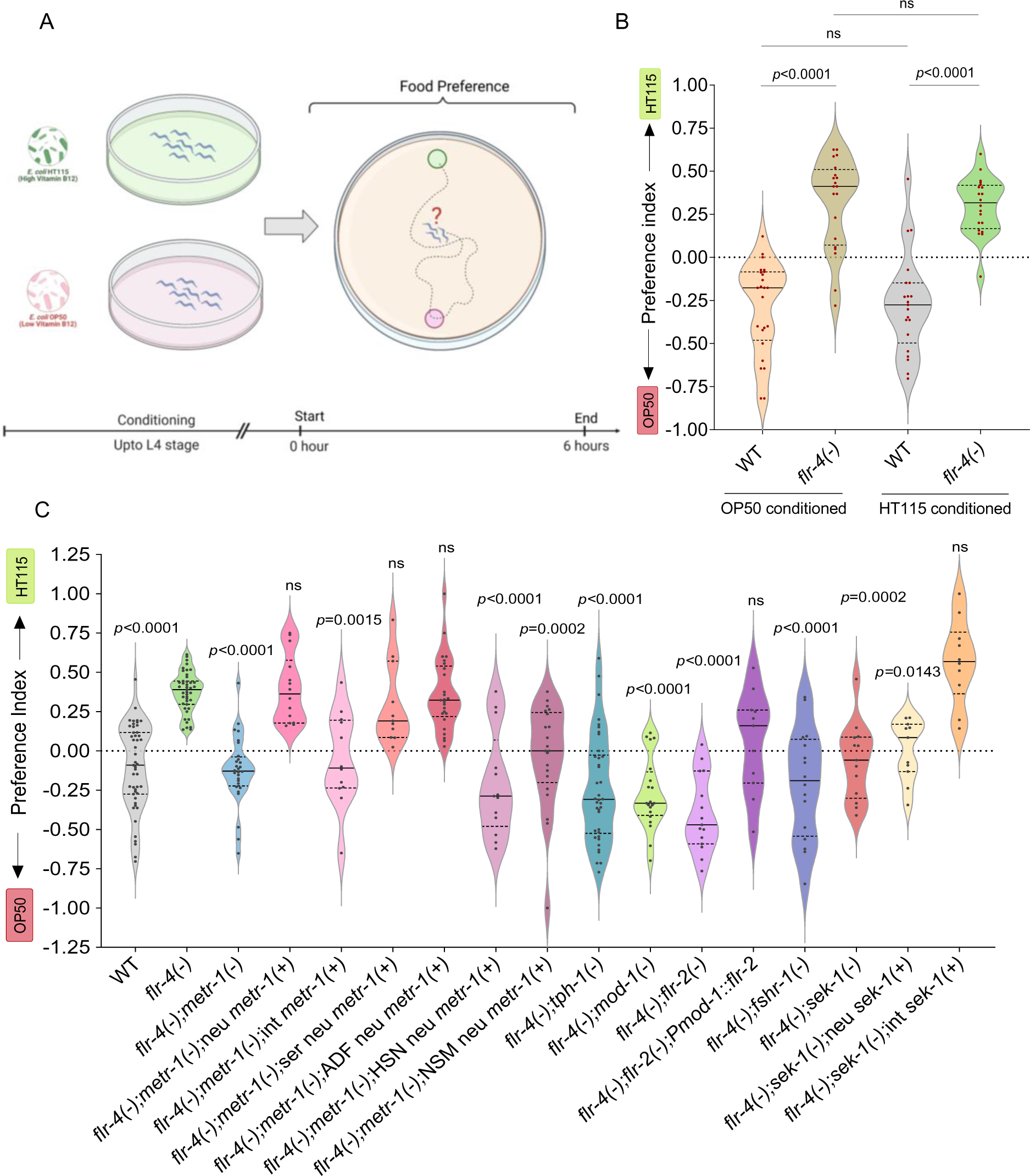
(A) Schematic of the food preference assay. (B) The *flr-4(n2259)* worms, upon conditioning with either OP50 or HT115 diet, displayed preference towards HT115, in contrast to wild-type worms which showed no significant preference towards either diet. Each dot represents one plate with over 100 worms. All groups were tested on at least three independent trials, in three separate biological replicates. *P*-value determined using Kruskal Wallis Test with Dunn’s multiple comparisons test. (C) The food preference of *flr-4(n2259)* worms towards HT115 diet was reversed when *metr-1, tph-1, mod-1, flr-2*, *fshr-1* or *sek-1* was mutated, as in *flr-4(n2259);metr-1(ok521)*, *flr-4(n2259);tph-1(mg280), flr-4(n2259);mod-1(ok103)*, *flr-4(n2259);flr-2(ut5), flr-4(n2259);fshr-1(ok778)* or *flr-4(n2259);sek-1(km4)*, respectively. Moreover, the rescue of *metr-1* in all neurons [neu *metr-1(+)*], serotonergic neurons [ser neu *metr-1(+)*] or ADF neurons [ADF neu *(metr-1(+)*], but not in HSN or NSM neurons of *flr-4(n2259);metr-1(ok521)* and intestinal rescue of *sek-1* [int *sek-1(+)*], but not the neuronal rescue, in *flr-4(n2259);sek-1(km4)* worms was sufficient to restore the altered food preference to that of *flr-4(n2259)* worms. Further, driving the expression of *flr-2* in the MOD-1-containing neurons rescued the food preference of *flr-4(n2259);flr-2(ut5)* worms. Each dot represents one plate with over 100 worms. All groups were tested on at least three independent trials, in three separate biological replicates. *P*-value determined using Kruskal Wallis Test with Dunn’s multiple comparisons test, in comparison with *flr-4(n2259).* All experiments were performed at 20°C. All data and analysis are provided in the Source Data file.

**Figure 8.**
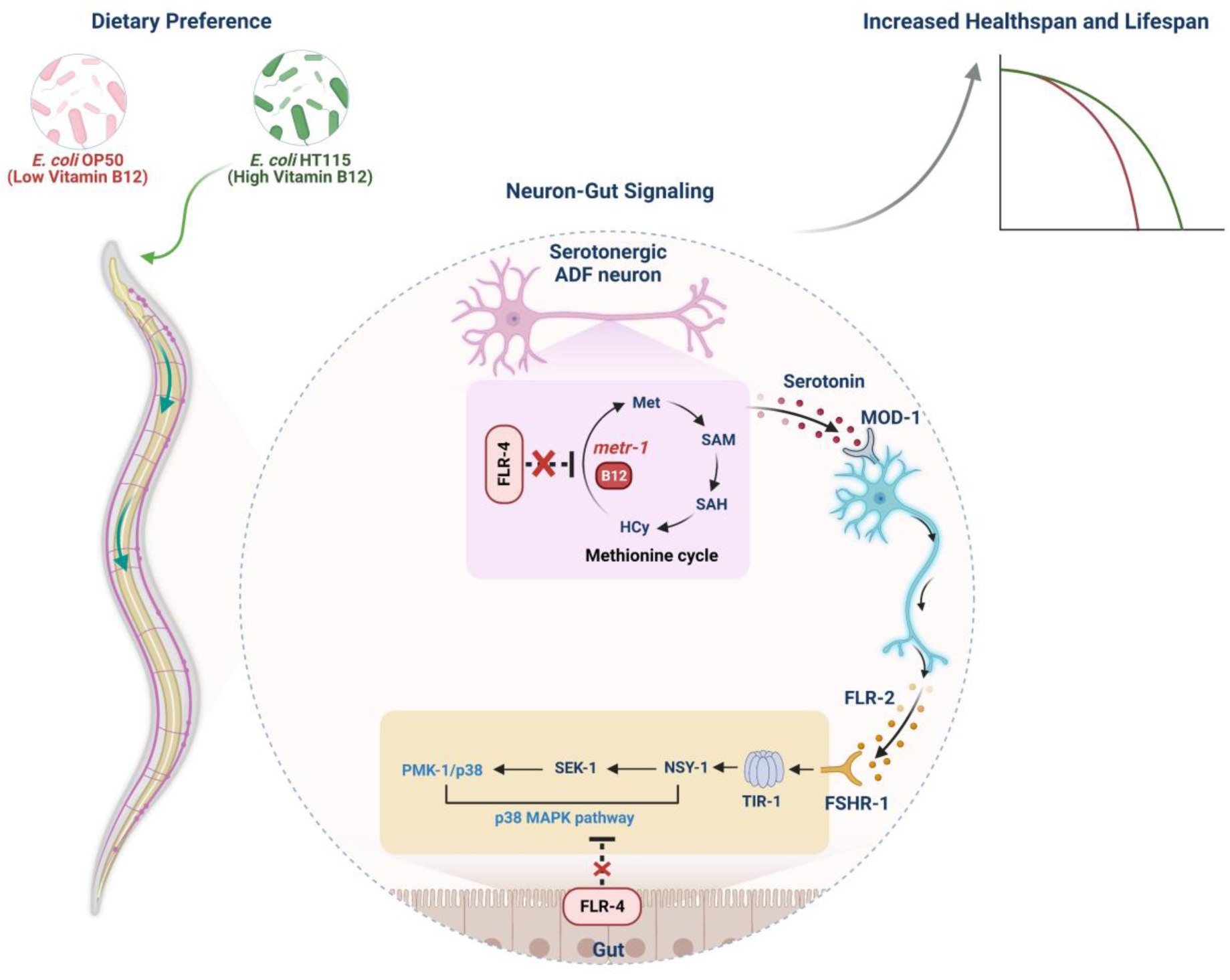
A model delineating the possible mechanism by which *C. elegans* FLR-4 modulates longevity through a neuron-gut crosstalk, preventing aberrant activation of the Met-C and the p38-MAPK pathway on diet containing different B12 levels. In the *flr-4* mutant, grown on a high B12 diet, the flux through the Met-C localized in the ADF serotonergic neurons directs the release of serotonin which binds to MOD-1, its cognate receptor in the interneurons. These interneurons, in turn, release the neuropeptide FLR-2 which then binds to its cognate receptor FSHR-1 in the intestine. This cascade induces the oligomerization of TIR-1, activating the downstream p38-MAPK pathway. Consequently, this leads to an increase in CyTP gene expression, enhancement of osmotic stress tolerance and an extension of lifespan of the *flr-4* mutant grown on high B12 diet.

Subsequently, we asked whether the foraging behavior towards the high B12 diet is also mediated by the reprogramming of the neuronal Met-C that activates the serotonin-FLR-2/FSHR-1-p38-MAPK axis. We observed that the preference of *flr-4(-)* worms towards HT115 is reversed when any of the components within the aforementioned cascade was mutated (**Figure 7C**) while the preference of the wild-type worms remained unaffected (**Figure S7B**). Interestingly, rescuing the Met-C specifically within the ADF serotonergic neurons (but not in the HSN or NSM neurons) or the p38-MAPK pathway in the intestine (but not in the neurons) was sufficient for the preference of the *flr-4(-)* worms towards the HT115 diet (**Figure 7C**). These experiments suggest that the foraging behavioral of the *flr-4(-)* mutant requires the same neuron-gut signaling regulated by Met-C in the neurons to maximize longevity benefits associated with a diet rich in B12 levels.

## Discussion

In this study, we elucidate an elegant neuron-gut axis that maintains normal life history traits in response to a diet of differential B12 content. This adaptive capacity is maintained by the *C. elegans* FLR-4 that prevents ectopic activation of the Met-C in the ADF serotonergic neurons and the p38-MAPK in the intestine. B12 is an important micronutrient; deficiency as well as excess has adverse effects on life history traits (Bito and Watanabe 2016; Ding et al. 2015; Macneil and Walhout 2013; Yilmaz and Walhout 2014). While deficiency lowers fertility, delays development, lowers pathogen resistance and shortens life span, excess B12 in the diet can accelerate growth rate, unnecessarily increase fecundity, and shorten life span (Macneil and Walhout 2013; Yilmaz and Walhout 2014); all these changes may jeopardize species survival. This justifies the evolution of proteins like FLR-4 in nematodes to prevent abrupt changes in life history traits when the worms forage on a diet of differential nutritional content in its ecological niche. This gene also provides us with a unique paradigm to decipher how B12 influences a conserved metabolic pathway in the neurons to control the physiology of the entire body and regulate behaviour and longevity. Since the bacteria is both the food as well as the microbiota of the worms, this paradigm also provides us with a genetically trackable model to study interspecies interactions that may be conserved in higher organisms.

The contribution of neurons in the regulation of longevity was recognized over two decades ago (Alcedo and Kenyon 2004; Wolkow et al. 2000; Apfeld and Kenyon 1999). Neurons are responsible for detecting environmental cues as well as internal changes in energy balance to coordinate metabolic homeostasis (Riera and Dillin 2016). The chemosensory neurons have earlier been shown to play an important role in modulating longevity (Alcedo and Kenyon 2004). Mutants that have impaired sensory cilia formation, and as a consequence defective sensory perception, are often long-lived (Apfeld and Kenyon 1999). Even the increased lifespan by dietary restriction is mediated by the transcription factor SKN-1 acting only in the two ASI sensory neurons (Bishop and Guarente 2007). The thermosensory AFD neurons extend lifespan at warm temperatures by enhancing the DAF-9 sterol hormone signaling (Lee and Kenyon 2009). Interestingly, the chemosensory serotonergic ADF neurons are known to be activated by environmental cues and also modulate food-dependent behaviors (Sze et al. 2000; Sawin, Ranganathan, and Horvitz 2000). These pair of neurons are responsible for exhibiting learned aversive behaviors upon exposure to pathogenic bacteria by modulating immune responses (Zhang, Lu, and Bargmann 2005; Xie et al. 2013). Food-associated odors perceived by ADF neurons are responsible for regulating DR-dependent longevity (Zhang et al. 2021). However, the role of ADF neurons in modulating B12-derived behavioral and physiological responses has not been previously described. In our study, we have shown that exposure to a high B12 diet modulates the Met-C in the ADF neurons to alter gene expression, stress tolerance, and behaviour.

Neuroendocrine signaling often directs the function of regulatory molecules in distal tissues such as the gut to influence longevity. The cool-sensitive neurons IL1 extend lifespan through regulation of FOXO/DAF-16 function in the intestine at lower temperature (Zhang et al. 2018). Neuronal FOXO/DAF-16 can communicate with the intestinal FOXO/DAF-16 to ensure longevity (Uno et al. 2021). The AFD thermosensory neurons respond to warm temperatures by activating the HSF-1 transcription factor in the intestine using serotonin signaling (Tatum et al. 2015). Neuronal mitochondrial or ER proteotoxic stress signal is communicated to the intestine to mount an Unfolded Stress Response (UPR) that modulates longevity (Berendzen et al. 2016; Taylor and Dillin 2013). Consistent with the important role of neuron-gut signaling to modulate longevity, our findings exemplify how neurons process information related to B12 content of food by modulating Met-C in the ADF neurons, transmitting them to downstream interneurons to modulate the p38 MAPK in the distal intestine.

The gut-microbiota employs multiple channels of communication to influence the brain, behavior and life-history traits (Zhang et al. 2017). Bacterially-derived metabolites transmit signals to the brain conveying information about the pathogenicity or commensalism (Meisel and Kim 2014). The neuromodulator tyramine, produced by the *Providencia*, a commensal bacterium colonizing the gut, is converted to octopamine by the *C. elegans,* influencing the host’s aversive sensory response (O’Donnell et al. 2020). The enteric serotonergic NSM neuron detects food ingestion via ion channels DEL-3 and DEL-7 that are localized to the sensory terminals of the enteric neuron in the gut and are required for the feeding-associated neuronal activation, which leads to slowing of the locomotion while the animal feeds (Rhoades et al. 2019). Recently, B12-producing bacteria that colonize the gut has been shown to modulate neuronal excitatory cholinergic signaling and influence behavior (Kang et al. 2024). Our study contributes to the growing understanding of host-microbiota interactions where the B12-rich gut microbiota is capable of influencing the nervous system to augment 5-HT biosynthesis. This, in turn, triggers downstream neuropeptide signaling, ultimately culminating in the activation of the p38-MAPK immune pathway within the intestinal milieu. The ensuing activation of the immune signaling pathway in the gut intricately modulates gene expression, stress tolerance, behavioral patterns, and ultimately, longevity.

Distinct gut microbiota profiles are observed in depressed individuals compared to healthy controls (Liang, Wu, and Jin 2018). Alterations in the composition of the human gut microbiota have been linked to mood disorders and neuropsychiatric conditions, with associated disruptions in neurotransmitter balance (Huang et al. 2019). Discernible changes in the metabolic, immune, and endocrine systems also connect the pathophysiology of depression to the gut microbiota (Caspani et al. 2019). Individuals afflicted with psychotic illnesses frequently exhibit deficiencies in folate and B12, contributing to diminished levels of S-adenosylmethionine (SAMe) in the cerebrospinal fluid (CSF) (Sharma et al. 2017). To overcome this, patients are often co-administered with antidepressant drugs (like Selective Serotonin Reuptake Inhibitors; SSRIs) and B12 supplements (Syed, Wasay, and Awan 2013; Coppen and Bolander-Gouaille 2005), although the mechanistic underpinnings remain elusive. Like humans, worms are entirely dependent on diet or microbiota for B12 supply (Watanabe and Bito 2018). Our investigation using the worm provides a useful paradigm where one can study mechanistic underpinnings of B12-mediated effects on serotonin signaling that affects behaviour.

## KEY RESOURCES TABLE

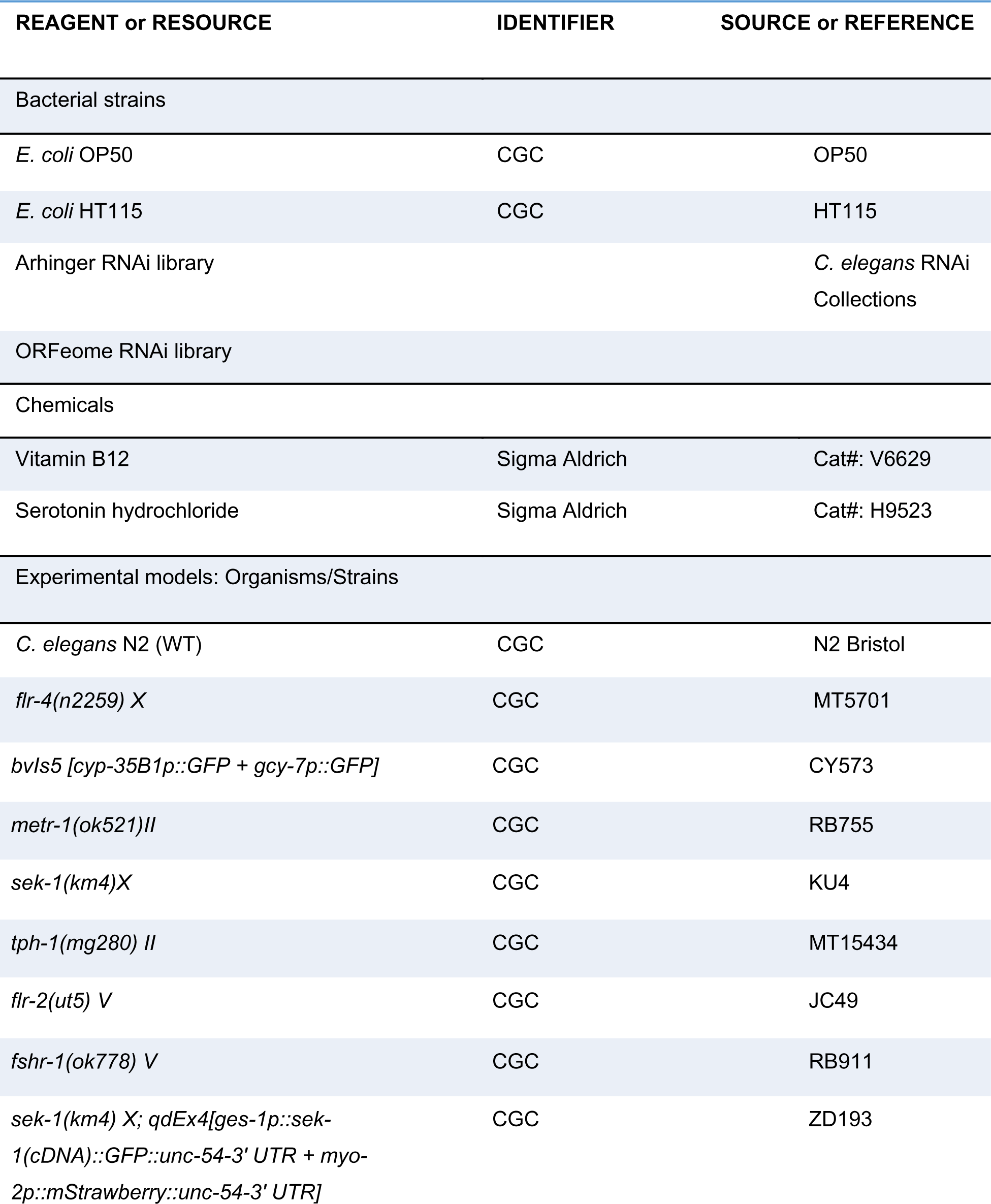
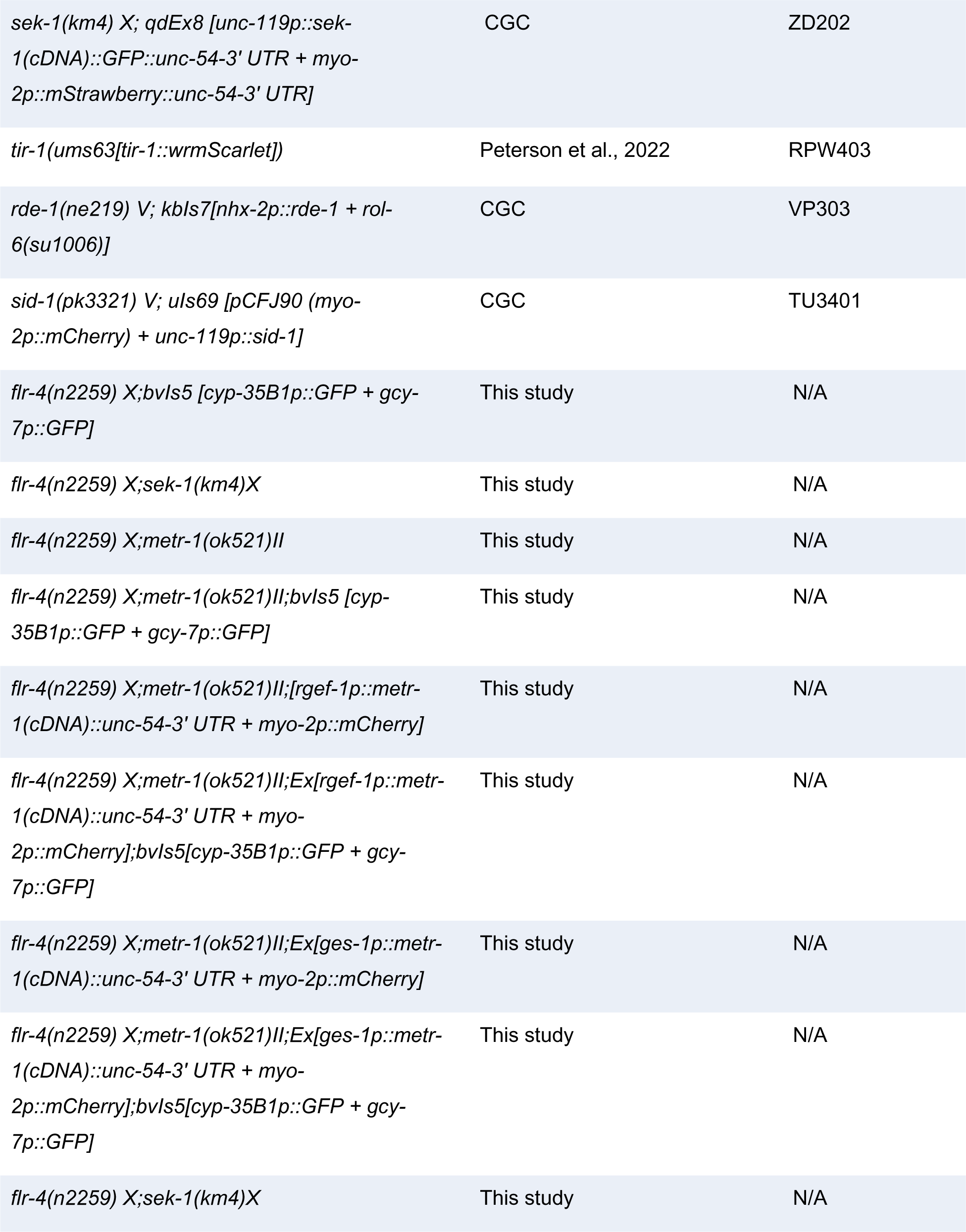
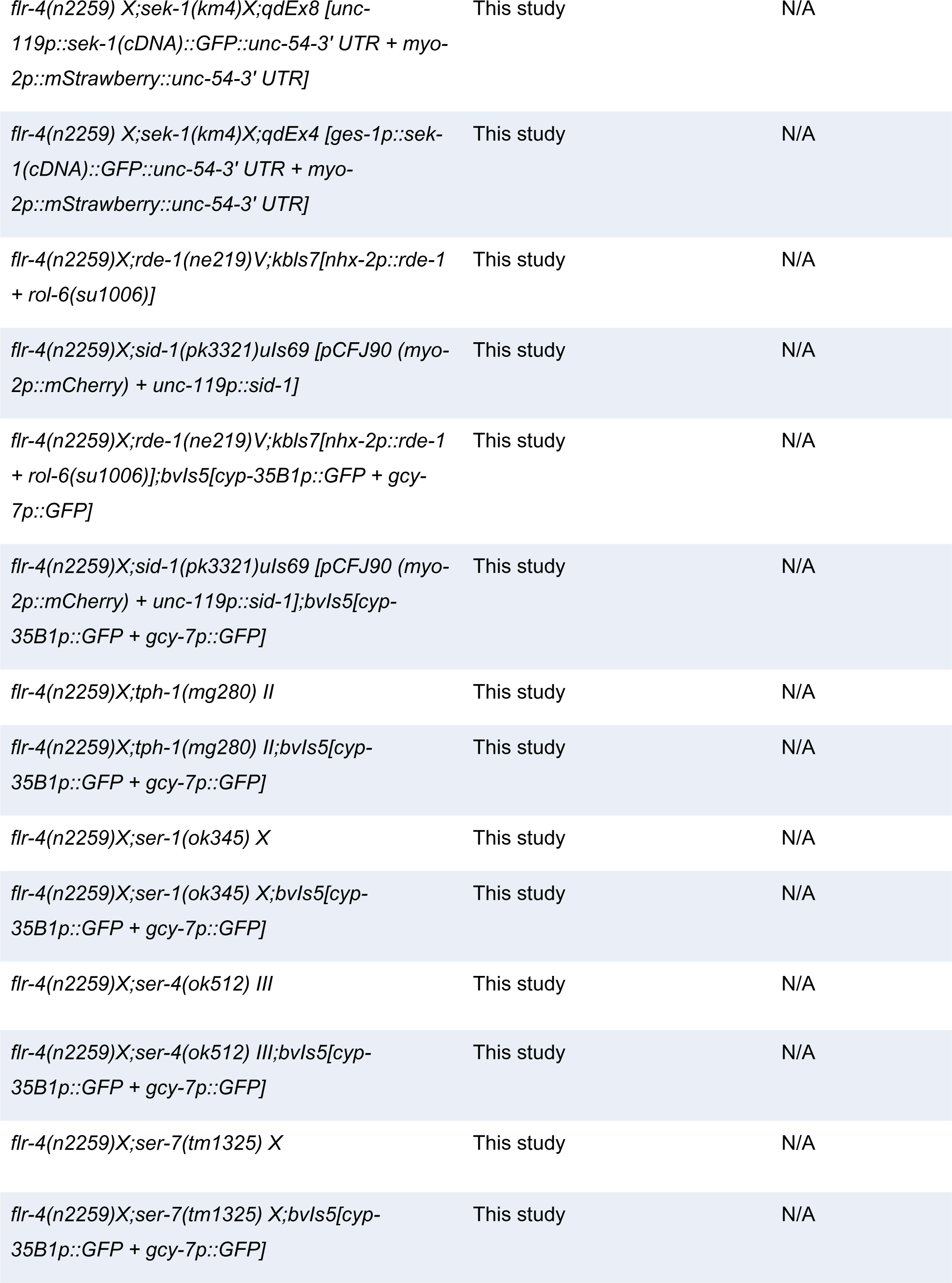
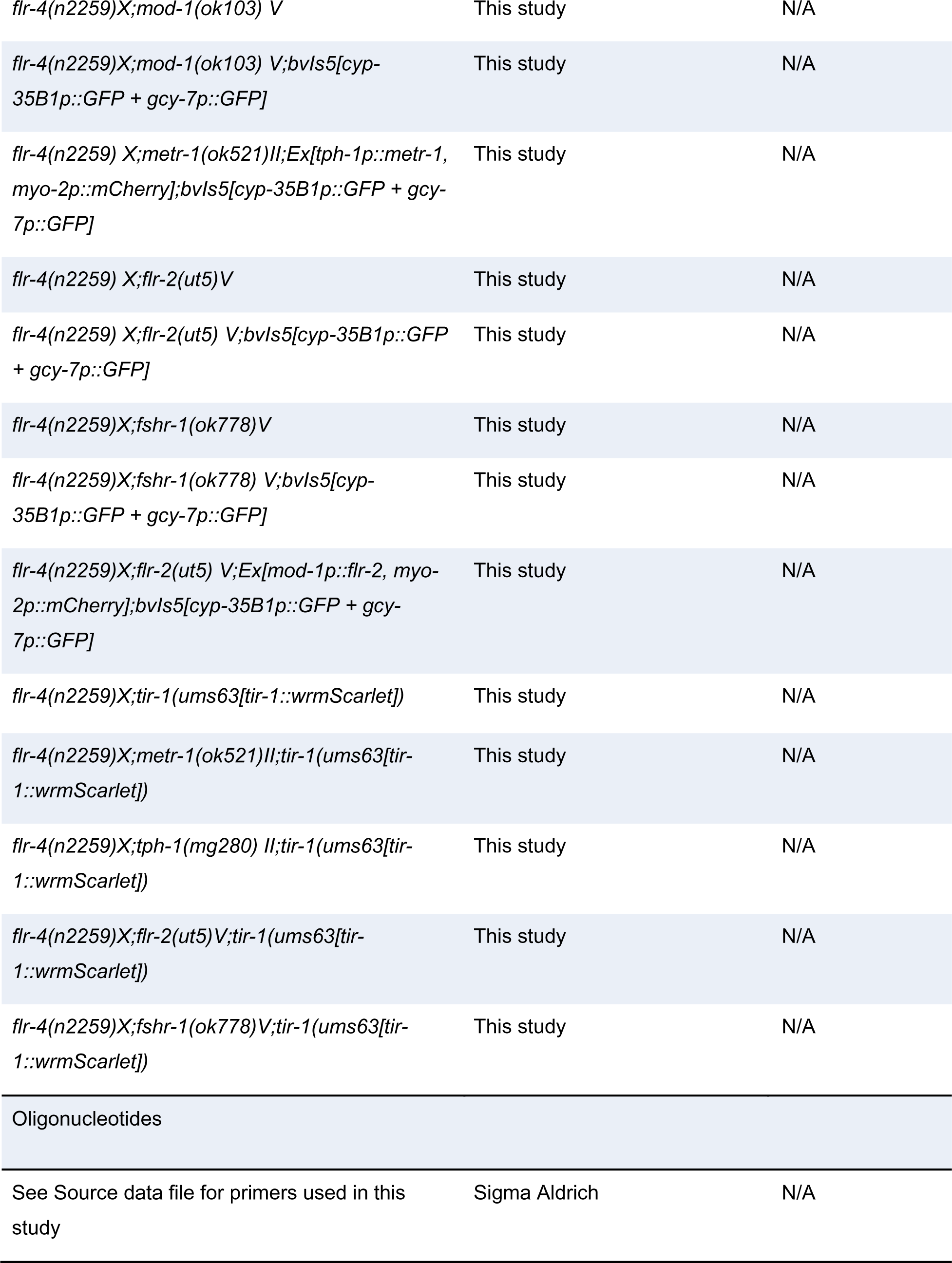
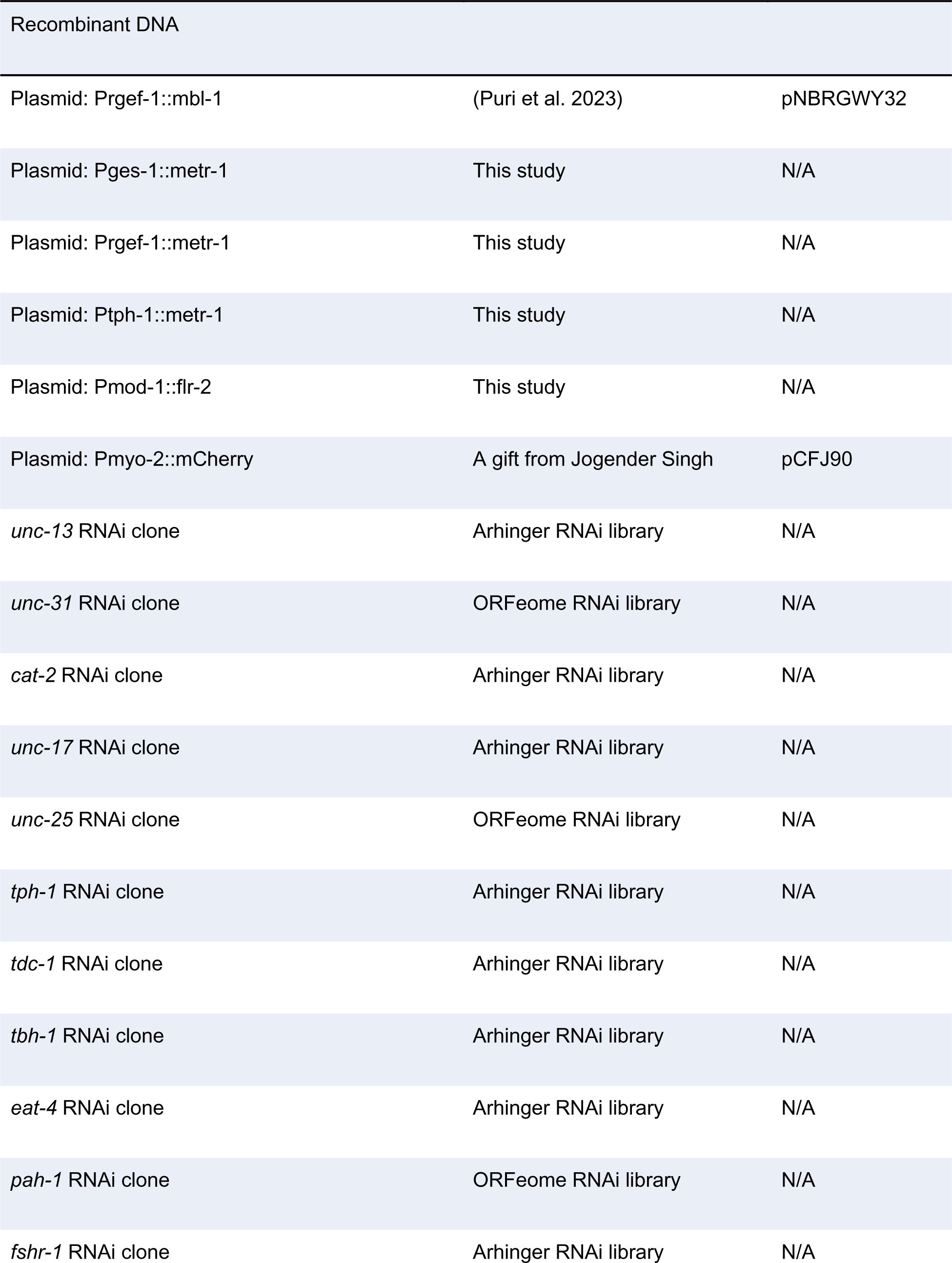
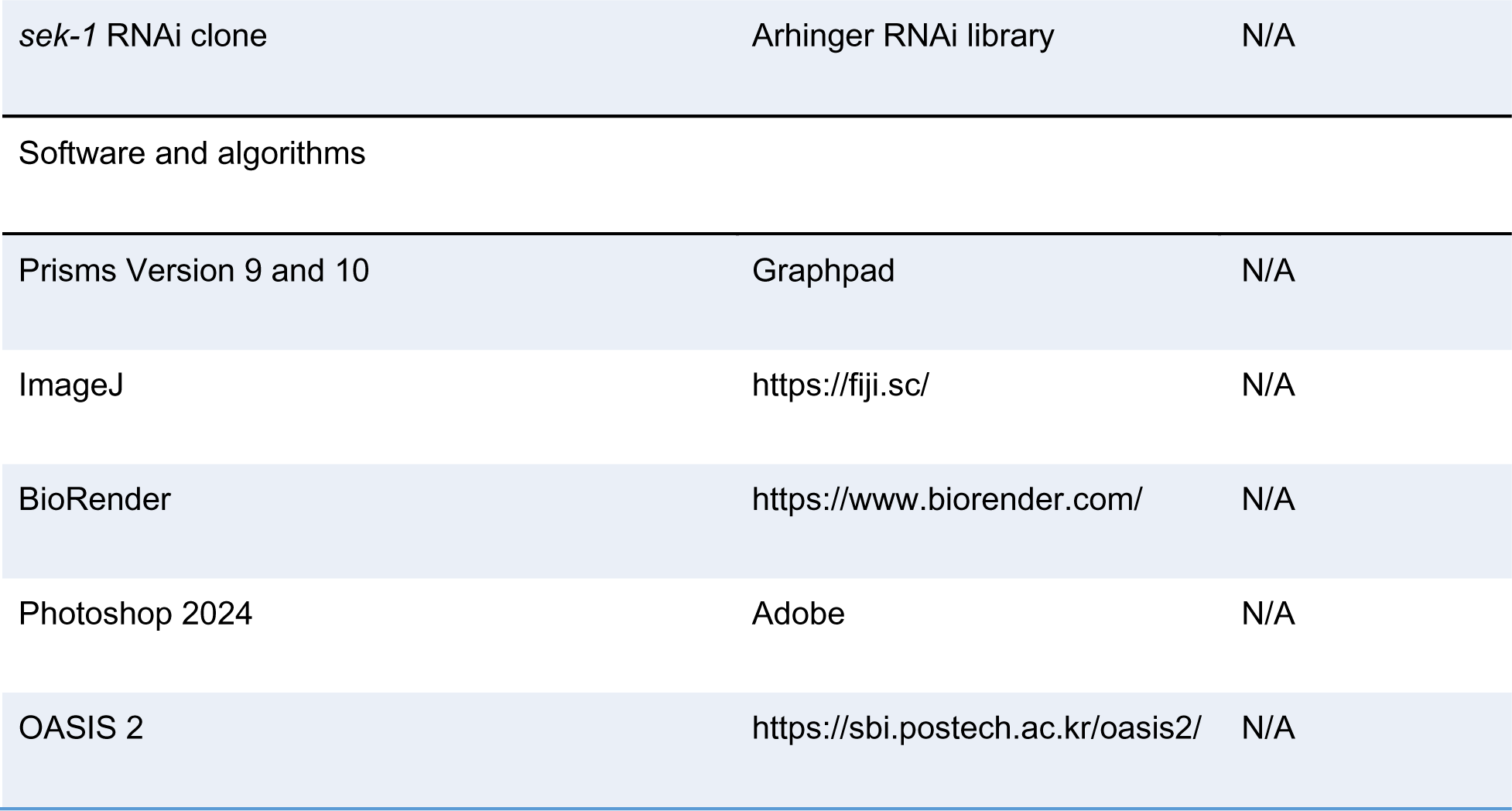

## RESOURCE AVAILABILITY

### Lead contact

Further information and requests for resources and reagents should be directed to and will be fulfilled by the lead contact, Arnab Mukhopadhyay.

### Materials availability

DNA constructs and *C. elegans* strains are available through the lead contact.

### *C elegans* growth and maintenance

All *C elegans* strains were maintained at 20°C on the Nematode Growth Medium (NGM) agar plates seeded with *Escherichia coli* OP50 bacteria as a food source for general maintenance. Depending upon the experiments, worms were either transferred to *Escherichia coli* OP50 or *Escherichia coli* HT115 seeded plates, post L1 synchronization by bleaching. Transgenic animals were constructed by microinjection. The Strains that were used in this study are provided in the key resources table.

## METHOD DETAILS

### DNA constructs and generation of transgenics

To clone *metr-1*(cDNA) under pan-neuronal promoter *Prgef-1,* plasmid pNBRGWY32 (*Prgef-1::mbl-1*) **(Puri et al. 2023)** was used. The plasmid was subjected to linearization through PCR, strategically designed to exclude the *mbl-1*(cDNA). The PCR amplified *metr-1*(cDNA) sequence was then fused with the linearized plasmid carrying the *Prgef-1* promoter through InFusion cloning (Takara) methodology. The *Prgef-1::metr-1* (10 ng/μl) construct, co-injection marker *Pmyo-2::mCherry* (2 ng/μl) and L4440 empty vector (70 ng/μl) were injected into *flr-4(-);metr-1(-)* and *flr-4(-);metr-1(-);Pcyp-35-b1::gfp* worms.

To clone *metr-1*(cDNA) under intestinal promoter *Pges-1,* plasmid *Prgef-1::metr-1* was used. The plasmid was subjected to linearization through PCR, strategically designed to exclude the *Prgef-1*. The PCR amplified 1931 bp of *Pges-1* sequence was then fused with the linearized plasmid carrying the *metr-1*(cDNA), through InFusion cloning (Takara) methodology. The *Pges-1::metr-1* (10 ng/μl) construct, co-injection marker *Pmyo-2::mCherry* (2 ng/μl) and L4440 empty vector (70 ng/μl) were injected into *flr-4(-);metr-1(-)* and *flr-4(-);metr-1(-);Pcyp-35-b1::gfp* worms.

To clone *metr-1*(cDNA) under serotonergic neuronal promoter *Ptph-1,* plasmid *Prgef-1::metr-1* was used. The plasmid was subjected to linearization through PCR, strategically designed to exclude the *Prgef-1*. The PCR amplified *Ptph-1* sequence was then fused with the linearized plasmid carrying the *metr-1*(cDNA), through InFusion cloning (Takara) methodology. The *Ptph-1::metr-1* (30 ng/μl) construct, co-injection marker *Pmyo-2::mCherry* (2 ng/μl) and L4440 empty vector (60 ng/μl) were injected into *flr-4(-);metr-1(-)* and *flr-4(-);metr-1(-);Pcyp-35-b1::gfp* worms.

To clone *metr-1*(cDNA) under serotonergic NSM neuronal promoter *Ptph-1(short)* of 158bp, plasmid *Prgef-1::metr-1* was used. The plasmid was subjected to linearization through PCR, strategically designed to exclude the *Prgef-1*. The PCR amplified *Ptph-1(short)* sequence was then fused with the linearized plasmid carrying the *metr-1*(cDNA), through InFusion cloning (Takara) methodology. The *Ptph-1(short)::metr-1* (30 ng/μl) construct, co-injection marker *Pmyo-2::mCherry* (2 ng/μl) and L4440 empty vector (60 ng/μl) were injected into *flr-4(-);metr-1(-)* and *flr-4(-);metr-1(-);Pcyp-35-b1::gfp* worms.

To clone *metr-1*(cDNA) under ADF serotonergic neuronal promoter *Psrh-142,* plasmid *Prgef-1::metr-1* was used. The plasmid was subjected to linearization through PCR, strategically designed to exclude the *Prgef-1*. The PCR amplified *Psrh-142* sequence was then fused with the linearized plasmid carrying the *metr-1*(cDNA), through InFusion cloning (Takara) methodology. The *Psrh-142::metr-1* (30 ng/μl) construct, co-injection marker *Pmyo-2::mCherry* (2 ng/μl) and L4440 empty vector (60 ng/μl) were injected into *flr-4(-);metr-1(-)* and *flr-4(-);metr-1(-);Pcyp-35-b1::gfp* worms.

To clone *metr-1*(cDNA) under HSN serotonergic neuronal promoter *Pegl-6,* plasmid *Prgef-1::metr-1* was used. The plasmid was subjected to linearization through PCR, strategically designed to exclude the *Prgef-1*. The PCR amplified *Pegl-6* sequence was then fused with the linearized plasmid carrying the *metr-1*(cDNA), through InFusion cloning (Takara) methodology. The *Pegl-6::metr-1* (30 ng/μl) construct, co-injection marker *Pmyo-2::mCherry* (2 ng/μl) and L4440 empty vector (60 ng/μl) were injected into *flr-4(-);metr-1(-)* and *flr-4(-);metr-1(-);Pcyp-35-b1::gfp* worms.

To clone *flr-2*(cDNA) under serotonin receptor promoter *Pmod-1,* plasmid *Prgef-1::metr-1* was used. The plasmid was subjected to linearization through PCR, strategically designed to exclude the *Prgef-1*. The PCR amplified *Pmod-1* sequence was then fused with the linearized plasmid carrying the *mbl-1*(cDNA), through InFusion cloning (Takara) methodology. Subsequently, the PCR amplified *flr-2*(cDNA) was then fused with linearized *Pmod1::mbl1* plasmid, through InFusion cloning (Takara) methodology. The *Pmod-1::flr-2* (30 ng/μl) construct, co-injection marker *Pmyo-2::mCherry* (2 ng/μl) and L4440 empty vector (60 ng/μl) were injected into *flr-4(-);flr-2(-)* and *flr-4(-);flr-2(-);Pcyp-35-b1::gfp* worms.

### Bacterial growth

Bacteria stored in glycerol stocks were streaked onto Luria-Bertani (LB) plates and then allowed to incubate at 37°C for 16 hours until distinct single colonies became visible. A single colony was then used to inoculate LB broth and grown overnight at 37°C for the primary culture. The secondary cultures were initiated with a 1/100th volume of the primary culture and left to incubate at 37°C until the optical density (OD600) of 0.6 was attained. The bacterial pellets were then resuspended in 1/10th volume of 1XM9 buffer. Approximately 300 µl of bacterial suspension was spread on NGM RNAi plates and left to dry for 1 day. The following bacterial strains were utilized: *E. coli* OP50 and *E. coli* HT115

### RNA interference (RNAi) assay

The Ahringer or Vidal *C. elegans* RNAi-feeding library was utilized for obtaining RNAi-feeding bacteria. The RNAi plasmids were verified for the correct target sequence. NGM RNAi plates were prepared by supplementing with 100 µg/ml ampicillin and 4mM β-D-isothiogalactopyranoside (IPTG). RNAi bacteria were cultured overnight at 37°C in LB media supplemented with 100 µg/ml ampicillin and 12.5 µg/ml tetracycline for the primary culture. The secondary cultures were initiated with a 1/100th volume of the primary culture supplemented with 100 µg/ml ampicillin and left to incubate at 37°C until the OD600 of 0.6 was attained. The bacterial pellets were then resuspended in 1/10 th volume of 1XM9 buffer consisting of 100 µg/ml ampicillin and 1 mM IPTG. Approximately 300 µl of bacterial suspension was spread on NGM RNAi plates and left to dry for 1 day. Worms were exposed to RNAi plates from L1 onwards. For knocking down genes expressed in the neurons, late L4 stage worms were transferred to freshly seeded RNAi plates.

### Synchronization of worms

Gravid worms, cultivated on *E coli* OP50, were bleached, and the resulting eggs were exposed to L1 stage starvation for 16 hours in M9 buffer at 20°C. Subsequently, the L1 larvae were subjected to centrifugation at 2500 rpm for one minute, followed by aspiration of the buffer. The L1 larvae were then transferred to the designated experimental plates.

### Vitamin B12 supplementation assay

A 3mM stock solution of Vitamin B12 (V6629; Sigma, USA) was prepared in Milli-Q water. Subsequently, a working stock of 1 mM was prepared by diluting the 3 mM stock with 1XM9 buffer and then filtered. To the 10-fold concentrated secondary culture, a specified volume of 1µM Vitamin B12 was added, to make 64nM of Vitamin B12-supplemented bacterial feed.

### Serotonin supplementation assay

A 100mM stock solution of Serotonin hydrochloride (H9523; Sigma, USA) was prepared in a 1XM9 buffer. A specified volume of 100mM was added to NGM plates seeded with bacteria to make a final plate concentration of 5mM. Plates were allowed to dry for approximately 2 hours. Late L3 worms were then transferred on the serotonin-supplemented bacterial plates.

### Osmotic stress assay

The worms were raised until they reached the L4 stage on respective bacterial-seeded NGM plates, which contained 50 mM Sodium chloride (NaCl) and were seeded with various types of bacterial feeds (either supplemented or with RNAi). At the L4 stage, worms were then placed on unseeded NGM plates, which contained 350 mM NaCl, for 9 minutes. After this 9^th^ minute, the worms were gently transferred onto unseeded NGM plates with 50 mM NaCl. The proportion of worms exhibiting movement was assessed over a 15-minute timeframe to determine the percentage of recovery. For conducting statistical analyses on the recovery curve, P-values were determined by Two-way ANOVA using the GraphPad 10 software.

### Life span assay

L1 synchronized worms were grown on NGM plates seeded with respective bacterial strains. Upon reaching the late L4 stage, the worms were transferred to plates overlaid with 5-fluorodeoxyuridine (FUDR; final concentration of 100 µg/ml). Life span assessment was started on the 7^th^ day of adulthood and was continued every alternate day. Worms were scored as alive or dead by gently tapping with the platinum wire. *P*-values between survival curves were determined using Log-rank (Mantel-Cox) test through OASIS 2 software (https://sbi.postech.ac.kr/oasis2/).

### Food preference assay

Overnight grown bacterial primary cultures, were freshly inoculated in LB broth and were grown until OD600 of 0.6 was attained at 37°C. The bacterial pellets were then resuspended in 1/10th volume of 1XM9 buffer. Binary choice plates were made by seeding 10 µl of distinct bacterial suspensions at two opposite points on the 90mm NGM assay plates, followed by an incubation period of 16 hrs at 20°C. On the next day, L4-stage worms conditioned on specific bacterial diets or supplements were collected in 1XM9 buffer and washed thrice to remove bacteria. Approximately, 100 worms were then placed into the centre of the choice plate. After 6 hours, the plates were examined and scored to observe the spatial distribution of the worms. The preference index *I* was calculated as *I* = (N_B12_ - N_O_)/(Total number of worms), where N_B12_ is the number of worms in the B12-rich diet i.e. either HT115 or OP50 supplemented with 64nM B12, and N_O_ is the number of worms in the OP50 diet.

### CyTP reporter imaging and analysis

For Pcyp-35B1::gfp expression visualization, all Pcyp-35B1::gfp expressing Day 1 adult worms, grown on either different diet, supplementation or RNAi, were mounted on 2% agarose coated slides using 20mM Sodium azide. The GFP fluorescence emitted by the worms was captured using an AxioImager M2 microscope (Carl Zeiss, Germany) at a magnification of 10X. Individual images were stitched using Adobe Photoshop. GFP fluorescence was quantified by ImageJ software and subsequent statistical analyses were performed by using Graphpad 10 software.

### TIR-1::wrmScarlet puncta imaging and analysis

To visualize the expression of TIR-1::wrmScarlet, worms expressing TIR-1::wrmScarlet were subjected to *glo-3* RNAi from the L1 stage onwards, to deplete the autofluorescent gut granules **(Peterson et al. 2022).** For imaging L4 stage worms, were mounted on 2% agarose-coated slides using 20mM Sodium azide. At 63 X, using an LSM 980 confocal microscope (Carl Zeiss, Oberkochen, Germany), the last posterior intestinal region was identified under the DIC channel. Images were then captured under the red channel to visualize the specific TIR-1::wrmScarlet puncta and the green channel to observe the autofluorescent gut granules. All images were acquired in Z-stacks which were subsequently, compressed into a single image. Quantification of TIR-1::wrmScarlet puncta was executed using the ImageJ software. The region of interest (ROI) was selected using the free hand tool under the DIC channel. Subsequently, puncta in the red channel were identified using the Find Maxima tool, within the selected ROI. This information was then superimposed onto the green channel to identify the presence of autofluorescent puncta at the same spatial location. TIR-1::wrmScarlet puncta were considered as those exhibiting expression solely in the red channel and not in the green channel.

### Quantification and statistical analysis

All statistical methods used in the paper are described in the figure legends and additional details are provided in the method details. Statistics were computed using GraphPad Prism. All data and analysis are provided in the Source Data file.

## Acknowledgements

We thank the present and former members of the Molecular Aging laboratory (National Institute of Immunology) for their support. We are grateful to Dr. Anindya Ghosh Roy for his valuable suggestions and guidance, Drs. Manish Chamoli and Gautam Chandra Sarkar for critical reading of the manuscript. We thank Dr. Read Pukkila-Worley (University of Massachusetts Chan Medical School) for generously providing the *C. elegans* strain RPW403 *tir-1(ums63[tir-1::wrmScarlet])*. Some strains were provided by the *Caenorhabditis* Genetics Center (CGC), which is funded by the National Institutes of Health (NIH) Office of Research Infrastructure Programs (P40 OD010440).

## Competing interests

The authors declare no competing interests.

## Funding

This project was partly funded by the National Bioscience Award for Career Development (BT/HRD/NBA/38/04/2016), SERB-STAR award (STR/2019/000064), J.C. Bose Fellowship (JCB/2022/000021), SERB grant (CRG/2022/000525), DBT grant (BT/PR27603/GET/119/267/2018), and core funding from the National Institute of Immunology.

## Data availability statement

The data reported in this manuscript is available in the Source data file that accompanies the manuscript.

## Notes

### Competing Interest Statement

The authors have declared no competing interest.

