## Supplementary Figure for "Methionine cycle in a pair of serotonergic neurons regulates diet-dependent behavior and longevity through a neuron-gut signaling"

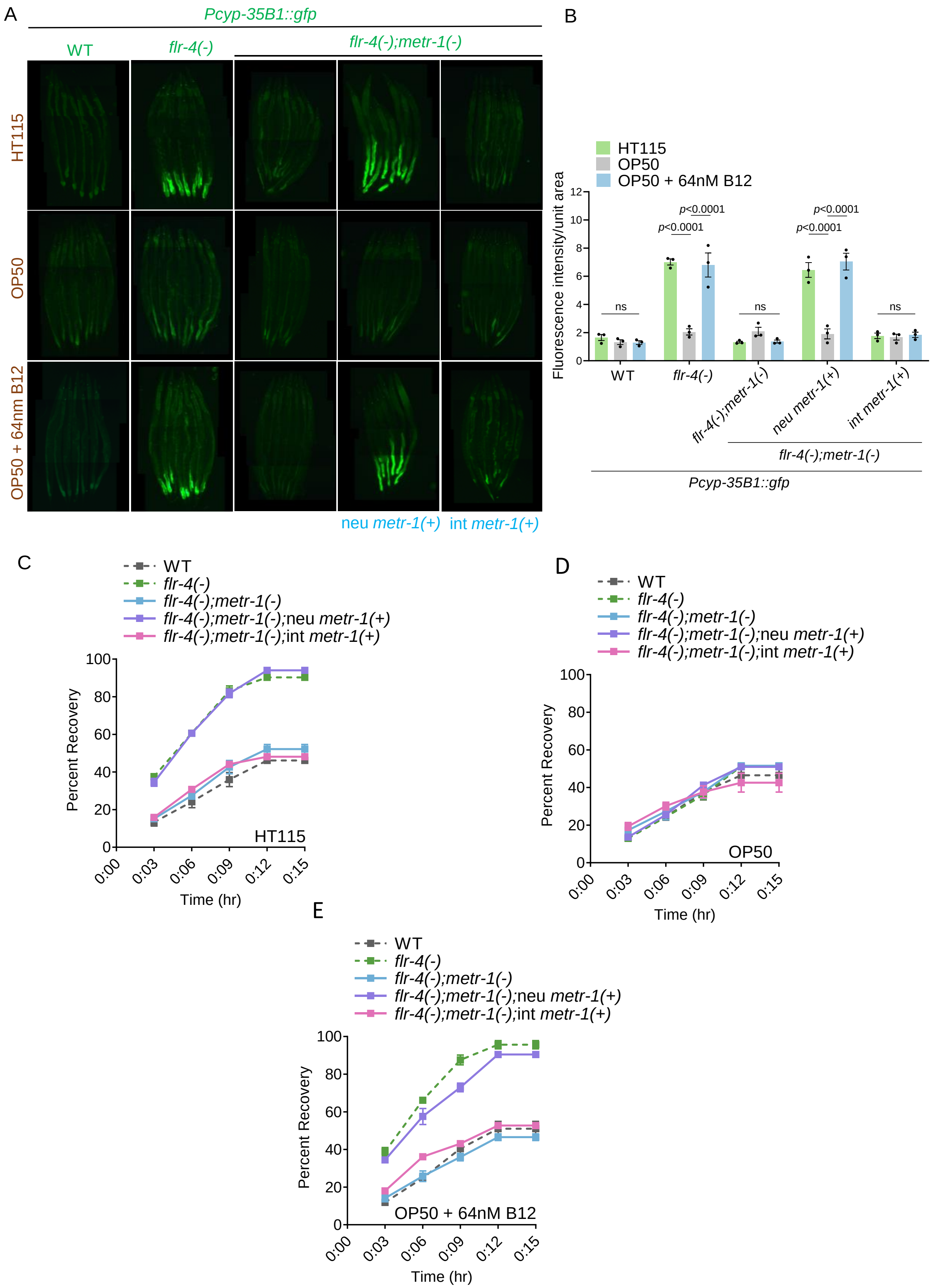
A
Pcyp-35B1::gfp
WT
flr-4(-)
flr-4(-);metr-1(-)
HT115
OP50
OP50 + 64nm B12
neu metr-1(+)
int metr-1(+)
B
HT115
OP50
OP50 + 64nM B12
Fluorescence intensity/unit area
ns
p<0.0001
p<0.0001
ns
p<0.0001
p<0.0001
ns
WT
flr-4(-)
flr-4(-);metr-1(-)
neu metr-1(+)
int metr-1(+)
flr-4(-);metr-1(-)
Pcyp-35B1::gfp
C
WT
flr-4(-)
flr-4(-);metr-1(-)
flr-4(-);metr-1(-);neu metr-1(+)
flr-4(-);metr-1(-);int metr-1(+)
Percent Recovery
HT115
Time (hr)
D
WT
flr-4(-)
flr-4(-);metr-1(-)
flr-4(-);metr-1(-);neu metr-1(+)
flr-4(-);metr-1(-);int metr-1(+)
Percent Recovery
OP50
Time (hr)
E
WT
flr-4(-)
flr-4(-);metr-1(-)
flr-4(-);metr-1(-);neu metr-1(+)
flr-4(-);metr-1(-);int metr-1(+)
Percent Recovery
OP50 + 64nM B12
Time (hr)

**Figure S1.**

(A) The expression of *gfp* in *flr-4(n2259);metr-1(ok521);Pcyp35B1::gfp* worms grown on OP50 supplemented with 64nM B12 was restored when *metr-1* was rescued only in the neurons (using the pan-neuronal *rgef-1* promoter) [neu *metr-1(+)*] but not when rescued in the intestine (using the *ges-1* promoter) [int *metr-1(+)*]. One of three biologically independent replicates is shown.

(B) Quantification of (A). Average of three biological replicates ± SEM. *P*-value determined using Two-way ANOVA with Tukey's multiple comparisons test.

(C-E) The osmotic stress tolerance of *flr-4(n2259);metr-1(ok521)* worms grown on HT115 (C), OP50 (D) or OP50 supplemented with 64nM B12 (E) was restored when *metr-1* was rescued only in the neurons (using the pan-neuronal *rgef-1* promoter) [neu *metr-1(+)*] in HT115 or OP50 supplemented with 64nM B12 diets, but not when rescued in the intestine (using the *ges-1* promoter) [int *metr-1(+)*]. One of three biologically independent replicates is shown.

Summary of osmotic stress tolerance assays are provided in the Source Data file. All experiments were performed at 20 $^{o}$C. All data and analysis are provided in the Source Data file.

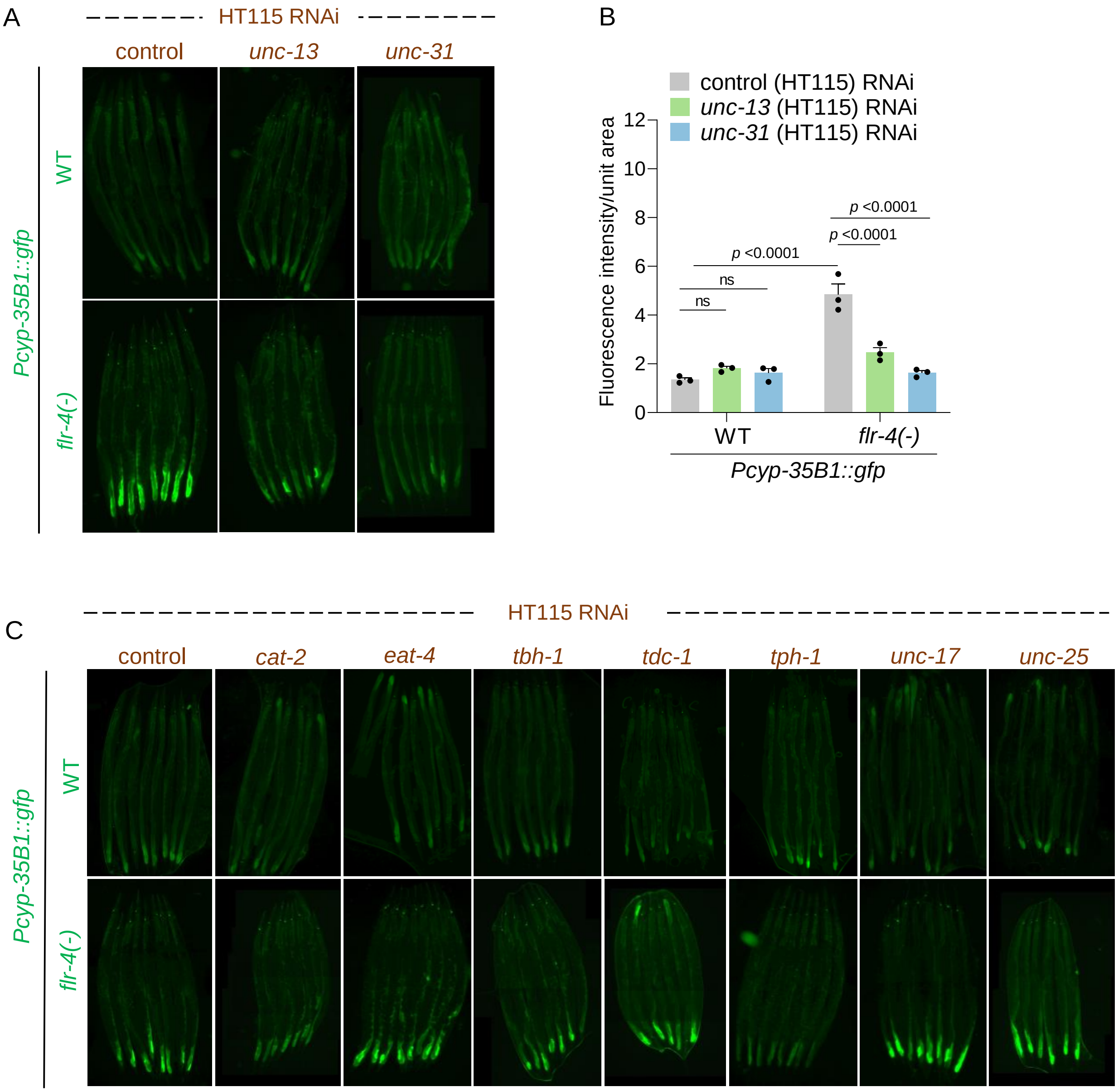
A
HT115 RNAi
control
unc-13
unc-31
Pcyp-35B1::gfp
WT
flr-4(-)
B
control (HT115) RNAi
unc-13 (HT115) RNAi
unc-31 (HT115) RNAi
Fluorescence intensity/unit area
0
2
4
6
8
10
12
ns
ns
p <0.0001
p <0.0001
p <0.0001
WT
flr-4(-)
Pcyp-35B1::gfp
C
HT115 RNAi
control
cat-2
eat-4
tbh-1
tdc-1
tph-1
unc-17
unc-25
Pcyp-35B1::gfp
WT
flr-4(-)

**Figure S2.**

(A) The expression of *gfp* in *flr-4(n2259);Pcyp35B1::gfp* worms were suppressed when genes involved in neurotransmitter release *(unc-13)* or neuropeptide release *(unc-31)* were knocked down using RNAi. One of three biologically independent replicates is shown.

(B) Quantification of (A). Average of three biological replicates ± SEM. *P*-value determined using Two-way ANOVA with Tukey's multiple comparisons test.

(C) The expression of *gfp* in *flr-4(n2259);Pcyp35B1::gfp* worms when genes involved in different neurotransmitter biosynthesis were knocked down by RNAi. One of three biologically independent replicates is shown.

All experiments were performed at 20 $^{o}$C. All data and analysis are provided in the Source Data file.

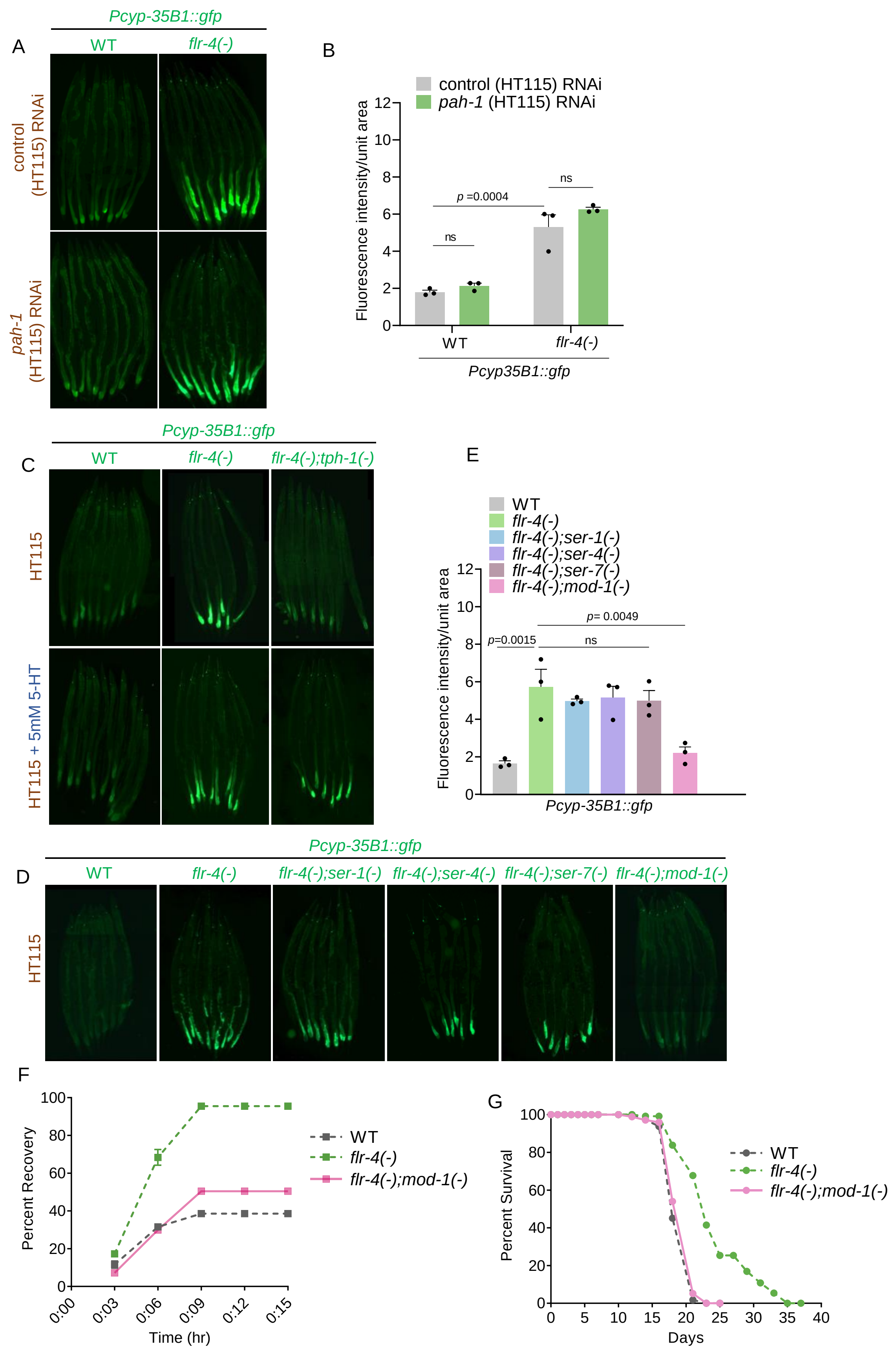
A
Pcyp-35B1::gfp
WT
flr-4(-)
control (HT115) RNAi
pah-1 (HT115) RNAi
B
control (HT115) RNAi
pah-1 (HT115) RNAi
Fluorescence intensity/unit area
p =0.0004
ns
ns
WT
flr-4(-)
Pcyp35B1::gfp
C
Pcyp-35B1::gfp
WT
flr-4(-)
flr-4(-);tph-1(-)
HT115
HT115 + 5mM 5-HT
E
WT
flr-4(-)
flr-4(-);ser-1(-)
flr-4(-);ser-4(-)
flr-4(-);ser-7(-)
flr-4(-);mod-1(-)
Fluorescence intensity/unit area
p=0.0015
p= 0.0049
ns
Pcyp-35B1::gfp
D
Pcyp-35B1::gfp
WT
flr-4(-)
flr-4(-);ser-1(-)
flr-4(-);ser-4(-)
flr-4(-);ser-7(-)
flr-4(-);mod-1(-)
HT115
F
Percent Recovery
Time (hr)
WT
flr-4(-)
flr-4(-);mod-1(-)
G
Percent Survival
Days
WT
flr-4(-)
flr-4(-);mod-1(-)

58

**Figure S3.**

(A) The expression of *gfp* in *flr-4(n2259);Pcyp35B1::gfp* worms was unaffected when the intestinal 5-HT biosynthesis gene *(pah-1)* was knocked down by RNAi. One of three biologically independent replicates is shown.

(B) Quantification of (A). Average of three biological replicates ± SEM. *P*-value determined using Two-way ANOVA with Tukey's Multiple Comparisons Test.

(C) The expression of *gfp* in *Pcyp35B1::gfp* and *flr-4(n2259);Pcyp35B1::gfp* worms were unaffected when supplemented with 5mM 5-HT. One of two biologically independent replicates is shown.

(D) The expression of *gfp* was suppressed in *flr-4(n2259);mod-1(ok103);Pcyp35B1::gfp* worms but not in *flr-4(n2259);ser-1(ok345);Pcyp35B1::gfp*, *flr-4(n2259);ser-4(ok512);Pcyp35B1::gfp*, or *flr-4(n2259);ser-7(tm1325);Pcyp35B1::gfp* worms. One of three biologically independent replicates is shown.

(E) Quantification of (D). Average of three biological replicates ± SEM. *P*-value determined using One-way ANOVA with Tukey's Multiple Comparisons Test.

(F) The increased osmotic tolerance of *flr-4(n2259)* was suppressed when *mod-1(ok103)* was mutated as in *flr-4(n2259);mod-1(ok103)*. One of three biologically independent replicates is shown.

(G) The life span of *flr-4(n2259)* worms was suppressed when *mod-1* was mutated, as in *flr-4(n2259);mod-1(ok103)*. One of three biologically independent replicates is shown.

Summary of osmotic stress tolerance and life span assays are provided in the Source Data file. All experiments were performed at 20 $^{o}$C. All data and analysis are provided in the Source Data file.

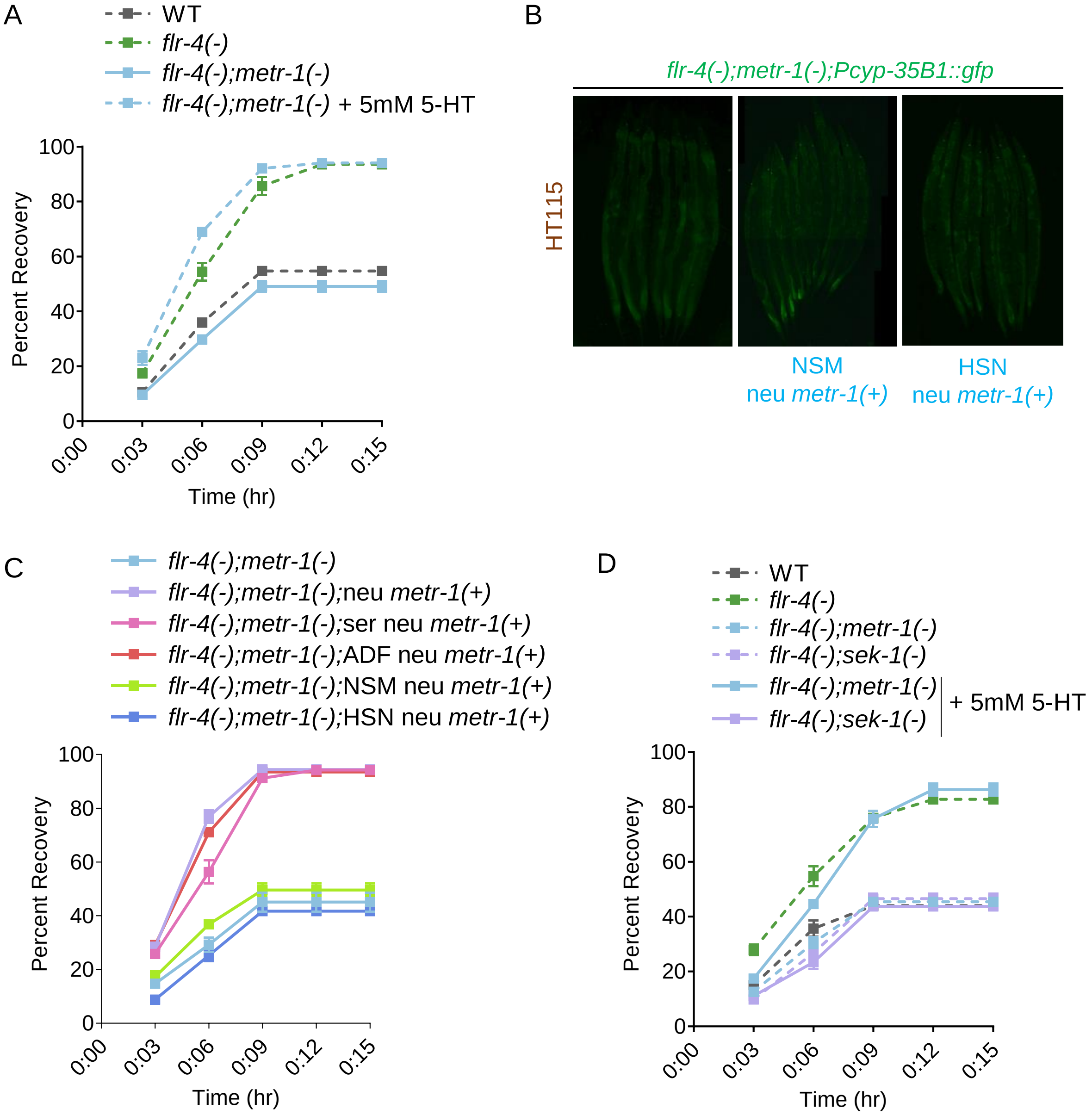
A
WT
flr-4(-)
flr-4(-);metr-1(-)
flr-4(-);metr-1(-) + 5mM 5-HT
Percent Recovery
Time (hr)
0:00
0:03
0:06
0:09
0:12
0:15
0
20
40
60
80
100
B
flr-4(-);metr-1(-);Pcyp-35B1::gfp
HT115
NSM
neu metr-1(+)
HSN
neu metr-1(+)
C
flr-4(-);metr-1(-)
flr-4(-);metr-1(-);neu metr-1(+)
flr-4(-);metr-1(-);ser neu metr-1(+)
flr-4(-);metr-1(-);ADF neu metr-1(+)
flr-4(-);metr-1(-);NSM neu metr-1(+)
flr-4(-);metr-1(-);HSN neu metr-1(+)
Percent Recovery
Time (hr)
D
WT
flr-4(-)
flr-4(-);metr-1(-)
flr-4(-);sek-1(-)
flr-4(-);metr-1(-)
flr-4(-);sek-1(-)
+ 5mM 5-HT
Percent Recovery
Time (hr)

**Figure S4.**

(A) Osmotic tolerance of *flr-4(n2259);metr-1(ok521)* was restored when supplemented with 5mM 5-HT. One of three biologically independent replicates is shown.

(B) The expression of *gfp* in *flr-4(n2259);metr-1(ok521);Pcyp35B1::gfp* worms was not restored when *metr-1* was rescued in the NSM serotonergic neurons (using the *tph-1* short promoter) [NSM *neu metr-1(+)*] or in the HSN serotonergic neurons (using the *egl-6* promoter) [HSN *neu metr-1(+)*]. One of three biologically independent replicates is shown.

(C) The osmotic stress tolerance of *flr-4(n2259);metr-1(ok521)* worms was restored when *metr-1* was rescued only in the serotonergic neurons (using the *tph-1* promoter) [*ser neu metr-1(+)*] or in the ADF serotonergic neurons (using the *srh-142* promoter) [ADF *neu metr-1(+)*]. One of two biologically independent replicates is shown.

(D) Osmotic tolerance of *flr-4(n2259);sek-1(km4)* was not restored when supplemented with 5mM 5-HT. One of three biologically independent replicates is shown.

Summary of osmotic stress tolerance are provided in the Source Data file. All experiments were performed at 20 $^{o}$C. All data and analysis are provided in the Source Data file.

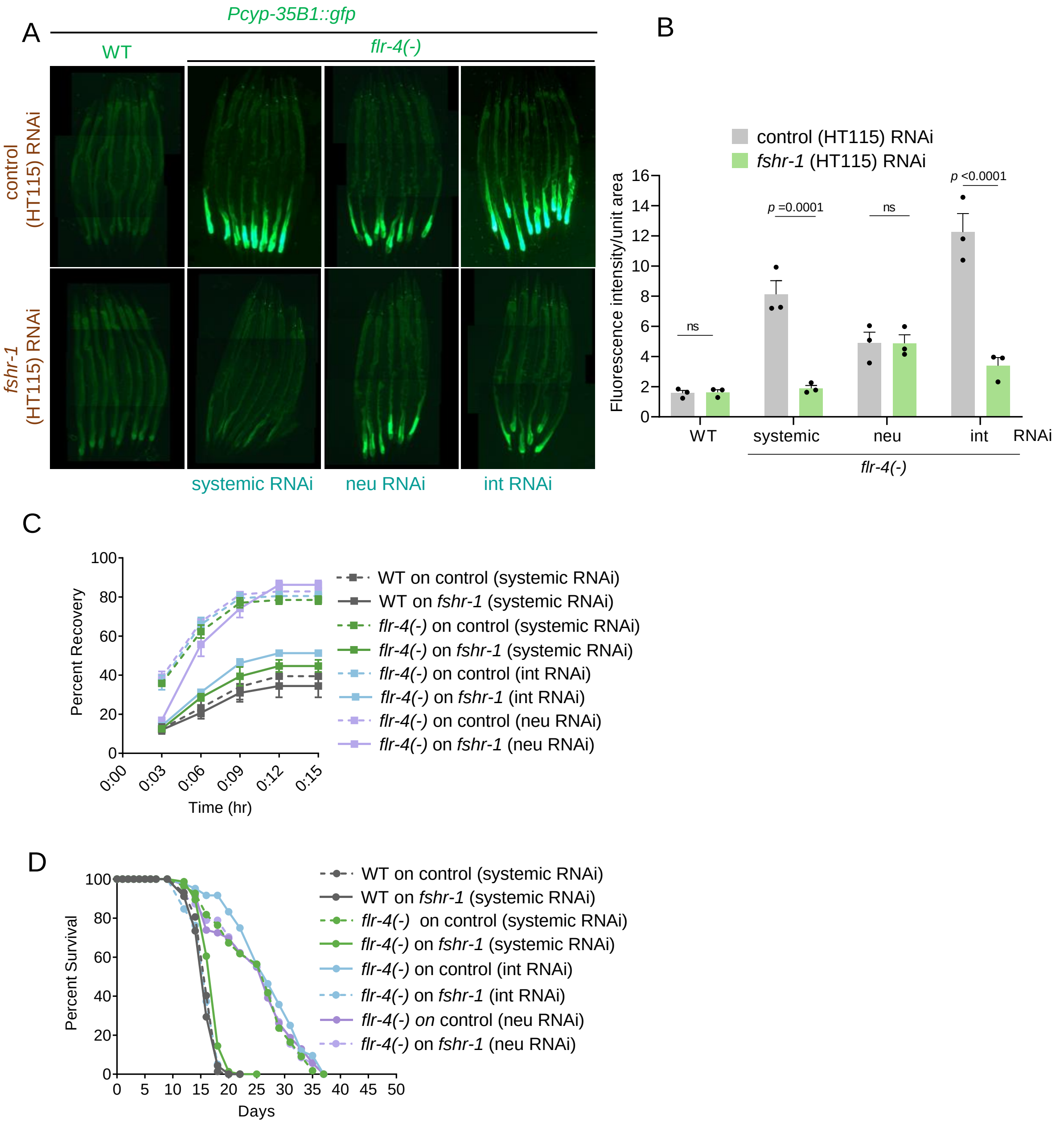
A
Pcyp-35B1::gfp
WT
flr-4(-)
control (HT115) RNAi
fshr-1 (HT115) RNAi
systemic RNAi
neu RNAi
int RNAi
B
control (HT115) RNAi
fshr-1 (HT115) RNAi
Fluorescence intensity/unit area
ns
p =0.0001
ns
p <0.0001
WT
systemic
neu
int
RNAi
flr-4(-)
C
Percent Recovery
Time (hr)
WT on control (systemic RNAi)
WT on fshr-1 (systemic RNAi)
flr-4(-) on control (systemic RNAi)
flr-4(-) on fshr-1 (systemic RNAi)
flr-4(-) on control (int RNAi)
flr-4(-) on fshr-1 (int RNAi)
flr-4(-) on control (neu RNAi)
flr-4(-) on fshr-1 (neu RNAi)
D
Percent Survival
Days
WT on control (systemic RNAi)
WT on fshr-1 (systemic RNAi)
flr-4(-) on control (systemic RNAi)
flr-4(-) on fshr-1 (systemic RNAi)
flr-4(-) on control (int RNAi)
flr-4(-) on fshr-1 (int RNAi)
flr-4(-) on control (neu RNAi)
flr-4(-) on fshr-1 (neu RNAi)

**Figure S5.**

(A) The expression of *gfp* was suppressed when *fshr-1* was knocked down in *flr-4(n2259); Pcyp35B1::gfp* (systemic RNAi) and *flr-4(n2259);rde-1(ne219);Pnhx-2::rde-1;Pcyp35B1::gfp* [intestine only RNAi (int RNAi)] but not in *flr-4(n2259);sid-1(pk3321);Punc-119::sid-1;Pcyp35B1::gfp* [neuron only RNAi (neu RNAi)] worms. One of three biologically independent replicates is shown.

(B) Quantification of (A). Average of three biological replicates ± SEM. *P*-value determined using Two-way ANOVA with Tukey's Multiple Comparison Test.

(C) Osmotic tolerance was suppressed when *fshr-1* was knocked down in *flr-4(n2259)* (systemic RNAi) and *flr-4(n2259);rde-1(ne219);Pnhx-2::rde-1* [intestine only RNAi (int RNAi)] but not in *flr-4(n2259);sid-1(pk3321);Punc-119::sid-1* [neuron only RNAi (neu RNAi)] worms. One of two biologically independent replicates is shown.

(D) The lifespan of *flr-4(n2259)* was suppressed when *fshr-1* was knocked down in *flr-4(n2259)* (systemic RNAi) and *flr-4(n2259);rde-1(ne219);Pnhx-2::rde-1* [intestine only RNAi (int RNAi)] but not in *flr-4(n2259);sid-1(pk3321);Punc-119::sid-1* [neuron only RNAi (neu RNAi)] worms. One of two biologically independent replicates is shown.

Summary of osmotic stress tolerance and life span assays are provided in the Source Data file. All experiments were performed at 20 $^{o}$C. All data and analysis are provided in the Source Data file.

A

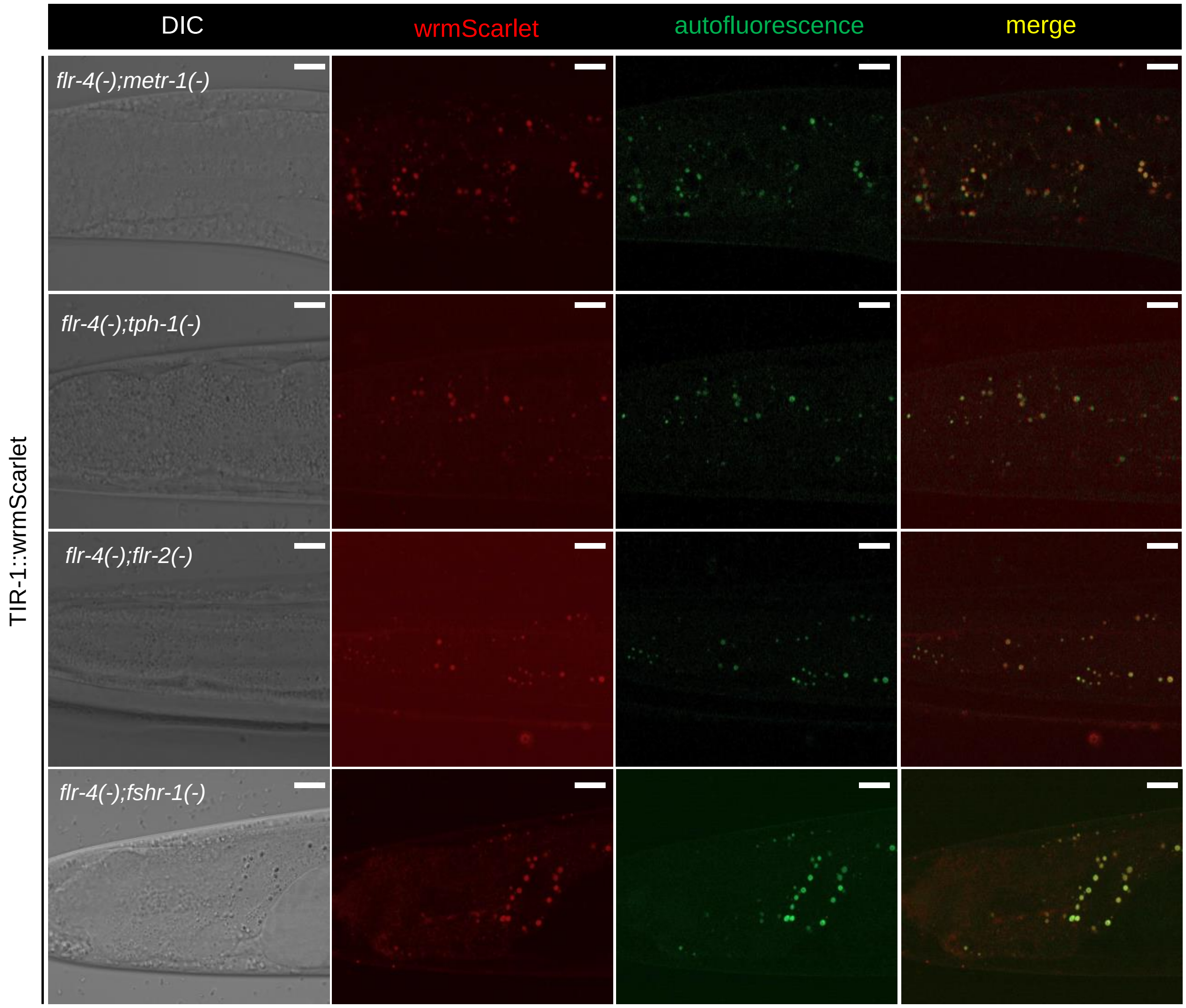
DIC
wrmScarlet
autofluorescence
merge
flr-4(-);metr-1(-)
flr-4(-);tph-1(-)
flr-4(-);flr-2(-)
flr-4(-);fshr-1(-)
TIR-1::wrmScarlet

**Figure S6.**

(A) The increased TIR-1 puncta formation in *flr-4(n2259)* worms was suppressed when *metr-1, tph-1, flr-2* or *fshr-1* was mutated, as in *flr-4(n2259);metr-1(ok521);tir-1::wrmScarlet, flr-4(n2259);tph-1(mg280);tir-1::wrmScarlet, flr-4(n2259);flr-2(ut5);tir-1::wrmScarlet* or *flr-4(n2259);fshr-1(ok778);tir-1::wrmScarlet*, respectively. One of three biologically independent replicates is shown.

All experiments were performed at 20 $^{o}$C. All data and analysis are provided in the Source Data file.

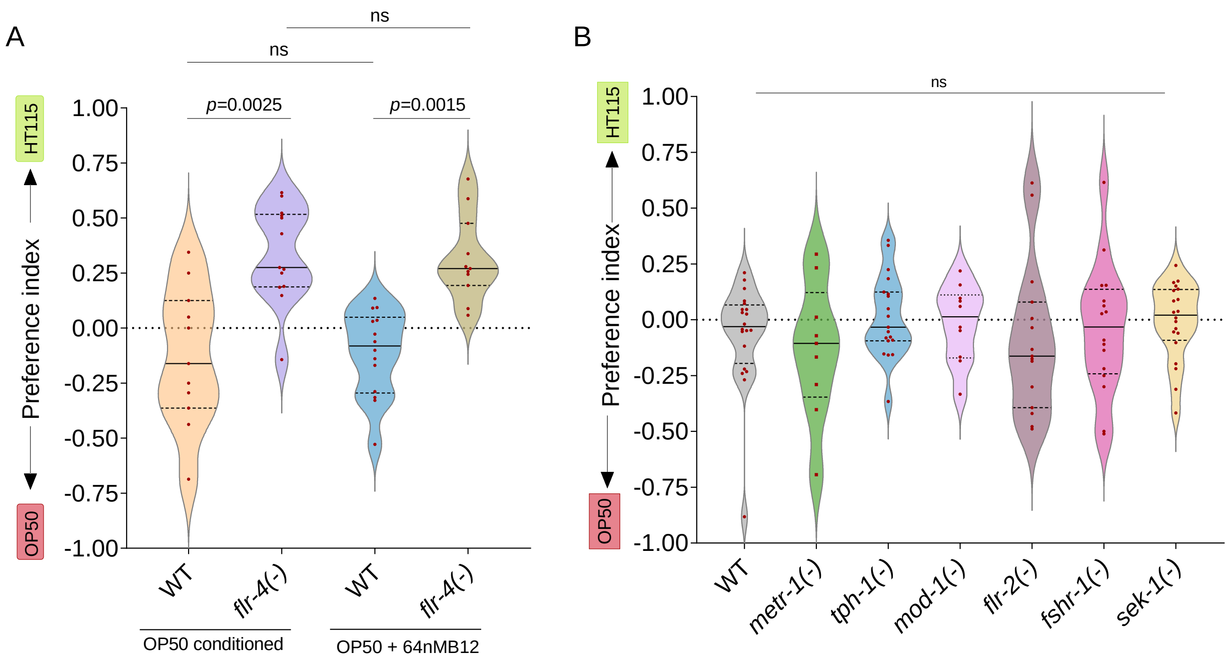

A
ns
ns
p=0.0025
p=0.0015
HT115
OP50
Preference index
1.00
0.75
0.50
0.25
0.00
-0.25
-0.50
-0.75
-1.00
WT
flr-4(-)
WT
flr-4(-)
OP50 conditioned
OP50 + 64nMB12
B
ns
HT115
OP50
Preference index
WT
metr-1(-)
tph-1(-)
mod-1(-)
flr-2(-)
fshr-1(-)
sek-1(-)

**Figure S7.**

(A) The *flr-4(n2259)* worms, upon conditioning with either OP50 or OP50 supplemented with 64nM B12 OP50 diet, display preference towards B12-rich diet, in contrast to wild-type worms which show no significant preference towards either diet. Each dot represents one plate with over 100 worms. All groups were tested on at least five independent trails, in three separate biological replicates. *P*-value determined using Kruskal Wallis Test with Dunn's multiple comparisons test.

(B) Wild-type worms as well as single mutants of *metr-1(ok521), sek-1(km4), tph-1(mg280), flr-2(ut5),* or *fshr-1(ok778)* displayed an absence of preference for either the OP50 or HT115 diet. Each dot represents one plate with over 100 worms. All groups were tested on at least three independent trails, in three separate biological replicates. *P*-value determined using Kruskal Wallis Test with Dunn's multiple comparisons test.

All experiments were performed at 20 $^{o}$C. All data and analysis are provided in the Source Data file.
